# Impact of Connectivity Granularity: A Comparison of ROI and Network-Level Approaches for Early Schizophrenia Classification

**DOI:** 10.1101/2025.09.30.679496

**Authors:** David Tomeček, Dominik Klepl, Zbyněk Pitra, Jiří Horáček, Jaroslav Tintěra, Filip Španiel, Jaroslav Hlinka

**Affiliations:** Early Episodes of SMI Research Center, National Institute of Mental Health, Czech Republic; Center for Advanced Studies of Brain and Consciousness, National Institute of Mental Health, Czech Republic; Department of Complex Systems, Institute of Computer Science, The Czech Academy of Sciences, Czech Republic; MR unit, Department of Diagnostic and Interventional Radiology, Institute for Clinical and Experimental Medicine, Czech Republic; Department of Psychiatry and Medical Psychology, Third Faculty of Medicine, Charles University, Czech Republic

## Abstract

While schizophrenia diagnosis relies on clinical interviews, there is growing interest in neuroimaging-based computational tools to aid classification. In particular, resting-state fMRI–derived functional connectivity has been explored as a potential biomarker. Here, we compare two common approaches to computing functional connectivity–region of interest (ROI)–level and brain network–level, and evaluate their predictive power for classifying first-episode schizophrenia patients, in contrast to most prior work focusing on chronic patients. The two approaches also differ in how many regions or components they use. We therefore compared them across a wide range of granularities: a functional Craddock parcellation computed at increasing sizes, from K = 10 to K = 950 regions, alongside the AAL90 atlas and ICA with 9, 17 and 80 components. The impact of granularity outweighed that of the parcellation method. Accuracy rose with granularity and plateaued at roughly 70 regions, with no further gain or loss at finer resolutions. Above the plateau the method mattered little: AAL90 was strongest (72.4% across classifiers, 79.6% for its best model) and an 80-component ICA matched it. Overall, our findings highlight region-based connectivity from a medium-resolution atlas as a promising representation for early-stage schizophrenia classification. These findings replicated when the trained classifiers were transferred to a dataset from another site.

## 1 Introduction

Schizophrenia is a severe mental disorder affecting ca. 1% of the population [Owen et al., 2011]. The abnormal developmental processes of the brain likely appear long before clinical symptoms of the disease [Stachowiak et al., 2013]. Based on the findings from the early neuroimaging studies, the disconnection hypothesis was proposed by Friston [1998] referring to reduced functional connectivity (FC). Generally, functional connectivity is defined as temporal correlation between spatially remote neurophysiological events [Friston et al., 1993, Friston and Frith, 1995]. The concept of disconnection (i.e., impaired connection) in schizophrenia was later described by Stephan et al. [2006] in terms of both structural and functional abnormal changes in the brain.

The common strategy for calculating functional connectivity relies on a priori-defined regions of interest (ROI) [Poldrack, 2007]. These regions of interest are commonly used in the form of atlases that parcellate the whole brain into several parts, e.g., the Talairach atlas [Talairach, 1988] or the automated anatomical labelling (AAL) atlas by Tzourio-Mazoyer et al. [2002]. Typically, an averaged time series is computed across the voxels covered by each region of the atlas, and FC is calculated between the averaged time series of the corresponding regions, reflecting their functional dependency [Poldrack, 2007]. The novel approaches enabled the study of the whole-brain connectivity patterns without the need to define an a priori ROI, namely the independent component analysis (ICA) [van den Heuvel and Pol, 2010]. ICA is a data-driven technique that can decompose the fMRI data into spatially independent components (in the case of spatial ICA) and their corresponding time series, where some of these components are related to sources of noise and several components are related to some known brain networks [Damoiseaux et al., 2006, Smith et al., 2009]. Functional network connectivity (FNC) is then calculated between the extracted components’ time series [Jafri et al., 2008]. However, the number of reported fMRI resting-state networks detected by the ICA decomposition is rather arbitrary and, in practice, depends on the amount of data available, choice of initial dimension reduction, and other factors. For instance, Beckmann et al. [2005] found eight brain networks in the resting-state fMRI data, similarly Shirer et al. [2012] identified 14 brain networks. On the other hand, Kiviniemi et al. [2009] reported 60 interpretable ICA components.

Several studies have been conducted describing impaired connectivity between multiple brain networks in schizophrenia (see Dong et al. [2018], or Ruiz-Torras et al. [2023]). However, due to methodological heterogeneity, the reported findings are highly inconsistent. The inconsistent results are likely partially due to variability in the selection of clinical samples as they include patients at different stages of the illness [Li et al., 2017]. Moreover, the studies are underpowered as they build their conclusions on small sample sizes with 40 participants on average [Dong et al., 2018]. Finally, methodological choices such as the selection of ROIs based on various brain parcellations might influence the results significantly.

Despite small sample sizes and heterogeneous clinical populations, machine learning approaches show promise in classifying schizophrenia based on FC-based features [Steardo et al., 2020, de Filippis et al., 2019]. Most proposed approaches utilise ROI-based connectivity features. While ICA-based functional network connectivity has been used less frequently, its initial application seemed to offer improved classification, particularly in chronic-stage classification offering up to 96% accuracy [Arbabshirani et al., 2013, 2014]. Overall, reported classification accuracies typically range from 74% to 93%, with higher performances more often reported using data recorded from chronic patients. On the other hand, classification of first-episode schizophrenia (FES) appears to be more challenging, likely due to the subtler or more variable neurobiological alterations early in the syndrome. Although some studies report high classification accuracy for FES cohorts, these studies are generally based on limited sample sizes, typically involving fewer than 100 subjects (in total over both groups) [Wang et al., 2018a, Liu et al., 2018, Zhuang et al., 2019], which restricts their generalisability.

A recent study evaluated support vector machine (SVM) classification potential on a dataset of 440 participants, systematically examining the impact of demographic subgroups (age and sex) and training sample size [Lee et al., 2022]. Accuracy increased gradually from 72.6% to 83.3% as sample size grew. Moreover, models trained on more diverse populations performed better than those restricted to a narrower age or sex subsample. These observations highlight the importance of considering both syndrome stage and dataset size when evaluating and comparing machine learning studies classifying schizophrenia.

Dadi et al. [2019] specifically addressed the variability in analytic pipelines for deriving functional connectome, highlighting key choices that influence classification performance. They emphasised the importance of functional data-driven region definitions, implemented as ICA decomposition with 80 networks. They also reported simple linear classifiers performing best across multiple populations and clinical settings.

Building on this work, we aim to contribute to the appreciation of the relative performance of inter-region and inter-network FC for classification of schizophrenia, and of how this performance depends on the granularity of the underlying parcellation.

Unlike previous work, we use a dataset of first-episode schizophrenia patients, a group that may pose a substantial challenge compared to chronic schizophrenia datasets often used in the literature [Dadi et al., 2019, Lee et al., 2022, Arbabshirani et al., 2013, 2014]. Using a resting-state fMRI dataset, we train several machine learning models to classify first-episode schizophrenia patients. For inter-network FC, we use the wide-spread approach of ICA; which reported very promising results in the study by Arbabshirani et al. [2013]. Although many competing methods for network definition exist, this is a commonly implemented and applied approach, representative in terms of performance of other network approaches [Dadi et al., 2019]. Specifically, we use various levels of granularity by including nine networks (as in the study by Arbabshirani et al. [2013]), 17 networks, and 80 networks (as suggested by Dadi et al. [2019], with the latter providing a finer subdivision that contrasts with the larger-scale networks typically discussed in resting-state studies).

For ROI-based FC, we use AAL [Tzourio-Mazoyer et al., 2002] as a commonly used anatomical atlas and the Craddock atlas [Craddock et al., 2012] at ten levels of granularity, from 10 to 950 regions. Comparing atlases to one another confounds granularity with how a region is defined. Varying the number of regions within a single scheme leaves granularity as the only quantity that changes, and the resulting curve serves as a reference for the other feature sets. If AAL90 and the ICA decompositions fall on it, their performance follows from their dimensionality alone; if they depart from it, the way their regions are defined contributes something beyond it.

## 2 Materials and methods

### 2.1 Participants

We obtained MRI data from 190 subjects in total (the primary cohort, hereafter referred to as IKEM) - 90 healthy controls (40 males), with an average age: 27.85, SD: 6.82, and 100 FES patients (58 males), with an average age: 28.79, SD: 6.84. FES patients were diagnosed according to ICD-10. The diagnostic procedure was standardised with a structured MINI International Neuropsychiatric Interview [Sheehan et al., 1998]. Patients were considered as FES if they fulfilled these criteria: a) first hospitalisation for schizophrenia, and b) clinical interview identified first psychotic and/or prodromal symptoms of psychosis not earlier than 24 months ago. The sample was checked for no significant difference in average age (*t* = *−*0.9382, *p* = 0.3493) and sex (*χ*^2^ = 2.9635, *p* = 0.0852) between the two groups. The exclusion criteria for groups included a history of seizures or significant head trauma, mental retardation, a history of substance dependence, and any MRI contraindications. The data set consists of resting-state fMRI data; the subjects were instructed to lie still with their eyes closed.

### 2.2 Data acquisition

The brain scans were acquired using a 3T Siemens Trio Tim MR scanner equipped with a standard 12-channel head coil at the Institute for Clinical and Experimental Medicine, Prague, the Czech Republic. Structural 3-dimensional (3D) images were obtained for anatomical reference using the T1-weighted (T1w) magnetization-prepared rapid gradient echo (MPRAGE) sequence with the following parameters: repetition time (TR) = 2300 ms, echo time (TE) = 4.6 ms, flip angle = 10°, voxel size = 1×1×1 mm^3^, field of view (FOV) = 256×256 mm, matrix size = 256×256, 224 sagittal slices. Functional images were obtained using the T2*-weighted (T2*w) gradient echo-planar imaging (GR-EPI) sequence sensitive to the blood oxygenation level-dependent (BOLD) signal with the following parameters: repetition time (TR) = 2000 ms, echo time (TE) = 30 ms, flip angle (FA) = 70°, voxel size = 3×3×3 mm^3^, field of view (FOV) = 144×192 mm, matrix size = 48×64, each volume with 35 axial slices (slice order: sequential decreasing), 400 volumes in total.

### 2.3 Data preprocessing

First, the structural and functional images were converted from DICOM to NIFTI format using the dcm2niix tool [Li et al., 2016]. The structural images were segmented into grey matter, white matter, and cerebrospinal fluid and directly normalised to MNI space. The following steps were performed with the functional data using a pipeline labelled as "default preprocessing pipeline for volume-based analyses (direct normalization to MNI-space)" in the CONN toolbox (https://web.conn-toolbox.org/) for Matlab 2016b (The MathWorks, Inc., Massachusetts, USA): functional realignment (motion correction) and unwarping, slice-timing correction, outlier identification, direct segmentation and normalisation, and functional smoothing with an 8-mm full width at half maximum (FWHM) kernel.

Following the preprocessing steps, we used two popular dimension reduction/data extraction techniques - ROI-based parcellation and spatial independent component analysis (ICA). In the first, ROI-based approach, we used the AAL atlas with 90 regions [Tzourio-Mazoyer et al., 2002] and the Craddock atlas [Craddock et al., 2012] in multiple versions, ranging from 10 to 950 regions, to examine how region granularity affects classification performance. The requested numbers of regions were 10, 30, 50, 70, 100, 200, 300, 400, 500 and 950; we refer to these as the ten granularity levels. The three finest levels (400, 500 and 950 regions) were analysed without lag, and, with the exception of the 400-region level, only in combination with PCA. The time series of each voxel was extracted and averaged across the corresponding region of each atlas. In the next step, we performed denoising of the extracted time series. Using linear regression, time series from each region were orthogonalised against five principal components of white matter (WM) signal, five principal components of cerebrospinal fluid (CSF) signal (WM and CSF masks were created from segmentation of the individual structural images), and six motion parameters with their temporal derivatives (translations and rotations in all three axes obtained when performing realignment (motion correction) of the functional images). The resulting time series were band-pass filtered with a window of 0.008–0.09 Hz and linearly detrended. For each subject, we then computed a functional connectivity (FC) matrix between the regions of each parcellation.

The EPI field of view did not cover every parcel in every participant. Seven subjects (six patients) exhibited grossly incomplete brain coverage and were excluded from all analyses and all feature sets, leaving 89 healthy controls and 94 patients. In the remaining subjects, individual small parcels of the finer Craddock levels still fell outside the field of view. Connectivity involving such a parcel is exported as a row and column of zeros rather than as missing data, and would therefore enter the classifiers as genuine, constant connectivity for the affected subjects alone. These parcels were identified from per-subject coverage tables and removed for every participant, taking the union over the two cohorts so that the primary and the validation datasets retain identical feature vectors. This leaves 198 of the 200 regions at the 200-region level and 296 of the 299 that the 300-region solution yields. At the three finest levels the discrepancy between the requested and the obtained number of parcels is larger: the Craddock solutions yield 392, 480 and 840 parcels rather than 400, 500 and 950, of which 381, 460 and 790 survive the coverage criterion. The coarser levels and AAL90 are unaffected. All region counts reported below, and the granularity axis of every figure, are these retained counts.

Every Craddock granularity level was supplied at its native resolution and resampled onto the functional grid internally by CONN when the regional time series were extracted, so that the atlas was treated identically at all ten levels.

In the second approach, we decomposed the functional data with spatial ICA, which was performed in the GIFT toolbox (https://trendscenter.org/software/gift/) for Matlab (The MathWorks, Inc., Massachusetts, USA). First, the data underwent a two-step reduction process using principal component analysis (PCA), and the minimum description length (MDL) principle was used to estimate the optimal number of independent components. For the ICA decomposition, we utilised the commonly used Infomax algorithm [Bell and Sejnowski, 1995]. After a visual inspection of the resulting 27 ICA components, ten of them were identified as artefactual and discarded. In this approach, we created a data set with a full set of 17 components (ICA17). Nine components, consistent with those found by Arbabshirani et al. [2013], were selected to form a second network-based dataset (ICA9). To ensure a fair comparison between network-based and ROI-based parcellations, we also performed an ICA decomposition with 80 components. This number was selected to approximate the dimensionality of commonly used ROI-based atlases and is in line with previous work [Dadi et al., 2019]. By aligning the number of components, we aimed to reduce the influence of differing feature dimensionality on model performance and isolate the effect of parcellation strategy. The extracted time series of the corresponding components were band-pass filtered with a window of 0.008–0.09 Hz and linearly detrended. Finally, we computed FNC between the selected components separately for the ICA9, ICA17, and ICA80 data sets. To prevent data leakage, the ICA decomposition was performed separately within each training fold, ensuring that test subjects were excluded from spatial map estimation and the resulting component time series were derived independently for the held-out fold.

As suggested by Arbabshirani et al. [2013], allowing lag between signals is important to account for variations in haemodynamic response shape among both brain regions and individuals. Therefore, we additionally prepared their lagged variants where the connectivity between pairs (regions or networks) corresponds to their maximum correlation from the interval from -3 to +3 seconds [Jafri et al., 2008, Arbabshirani et al., 2013]. In order to compute the lagged correlations within this interval with a one-second step, we linearly interpolated the original time series from each data set accordingly, as in the study by Arbabshirani et al. [2013].

The main comparison therefore rests on 14 parcellations - Craddock at 10, 30, 50, 70, 100, 200, 300, 400, 500 and 950 regions, AAL90, ICA9, ICA17 and ICA80 - each prepared in a non-lagged and a lagged variant, giving 28 feature sets in total.

The dimensionality of all feature sets except Craddock10, ICA9 and ICA17 exceeded the number of participants, so every feature set was additionally evaluated in a second configuration in which principal component analysis (PCA) is applied for dimensionality reduction, to rule out that the wider feature sets are limited simply by having more features than subjects.

The different sizes of the two tested subject groups (healthy controls and patients) could cause bias in the performance of classifiers in the next step. Therefore, five randomly selected subjects were removed from the patients’ group to balance the number of subjects in both classes (resulting in 89 healthy controls and 89 patients, 178 subjects in total). The reduced dataset was again checked for no significant difference in sex and age between the two groups. For comparability of results, all analyses were carried out on this reduced dataset.

### 2.4 Classification experiments

We trained 17 classifiers on each of the feature sets to classify healthy controls and first-episode schizophrenia patients: Linear Discriminant Analysis (LDA), Ridge Classifier (Ridge), Lasso, ElasticNet, Gaussian Naive Bayes (GaussianNB), k-Nearest Neighbours (KNN), Support Vector Machines with linear, RBF, sigmoid, and polynomial kernels (SVM-Linear, SVM-RBF, SVM-Sigmoid, SVM-Poly), Random Forest (RF), Extremely Randomised Trees (ExtraTrees), AdaBoost, Gradient Boosting (GradBoost), XGBoost [Chen and Guestrin, 2016], LightGBM [Ke et al., 2017], and a multilayer perceptron (MLP). The classifiers were selected to cover the range of approaches commonly applied to functional connectivity, spanning linear models, kernel methods, tree ensembles and neural networks.

Each classifier was evaluated on every feature set both with and without PCA, except at the 500- and 950-region levels, which were run with PCA alone due to the high computational cost. Every resulting configuration was tuned and evaluated by using a nested cross-validation.

To ensure reliable and unbiased performance estimates, we employed a nested cross-validation (nested CV) framework [Varoquaux et al., 2017, Varoquaux, 2018]. The outer cross-validation used five folds, which were fixed in advance and identical across all feature sets and classifiers, so that every classifier was evaluated on the same partitions. In the inner loop, hyperparameters were tuned independently for each outer fold by the tree-structured Parzen estimator, as implemented in Optuna [Akiba et al., 2019] and seeded identically for every fold, with 100 trials and stratified 5-fold cross-validation, maximising balanced accuracy. Optimised hyperparameters included classifier-specific regularisation strength, kernel parameters, and tree-related parameters where applicable. Inclusion of feature standardisation (zero mean, unit variance) was treated as a binary hyperparameter, allowing the inner CV to determine whether standardisation benefited each classifier–feature set combination. For classifiers using PCA to reduce the feature dimensionality, the number of retained components was similarly optimised via the inner CV, over the range from 2 to min(100*, p, n*_train_ *−* 1), where *p* denotes the number of features; in the submitted version it was instead fixed at 36, matching the number of connections in ICA9.

Classification performance was evaluated on the held-out test set of each outer fold using balanced accuracy as the primary metric, along with sensitivity, specificity, and the area under the receiver operating characteristic curve (AUC-ROC). Both cohorts are exactly class-balanced and every fold was stratified, so balanced accuracy coincides with overall accuracy; we refer to it simply as accuracy throughout. Final performance estimates are reported as means and standard deviations across the five outer folds.

Because most feature sets are wider than the sample, linear discriminant analysis used the singular-value solver throughout, which factorises the subjects-by-features matrix directly and yields a solution at every feature set. The eigenvalue solver, which requires a non-singular within-class scatter matrix, was excluded at all widths, and the shrinkage solver for parcellations of 200 regions or more, where its per-fit cost becomes prohibitive. All other feature sets, including AAL90 and ICA80, are unaffected.

### 2.5 External validation on an independent dataset

To test whether our findings generalised beyond the primary dataset, we transferred the classifiers trained and tuned on the IKEM cohort to an independent dataset acquired at a different site, without any re-training. Some loss of accuracy is therefore to be expected, and its size is itself one of the results. The independent dataset (NUDZ) comprised *N* = 174 subjects: 87 healthy controls (38 males), with an average age: 29.01, SD: 7.01, and 87 FES patients (38 males), with an average age: 28.95, SD: 6.93. We matched healthy controls to patients using a greedy nearest-neighbour algorithm that prioritised exact sex matching followed by minimal age difference. Additionally, we applied post-hoc filtering to remove sex-discordant pairs and those exceeding the Tukey outlier threshold for age differences (*Q*3+3*×IQR* = 3.79 years). The matched groups showed no significant differences in average age (*t* = 0.0564, *p* = 0.9551) or sex distribution (*χ*^2^ = 0.0, *p* = 1.0). For the validation dataset, we applied the same diagnostic as well as inclusion and exclusion criteria for the subjects as in the primary IKEM dataset. No data from the NUDZ dataset entered training, feature construction, or hyperparameter tuning. Because independent components are dataset-specific, we did not re-estimate ICA on the NUDZ dataset. Instead, each of the five ICA decompositions obtained from the IKEM training folds was applied to all NUDZ participants, and the corresponding tuned classifier was evaluated on them. Performance is reported as the mean (standard deviation) over the five IKEM-fold models.

## 3 Results

We selected three crucial modelling choices: the granularity of the brain parcellation, method of FC estimation, and inclusion of dimensionality reduction, to test their influence on the classification of first-episode schizophrenia patients. We compare the overall effects that these three choices have on classification (Fig. 1; Table 1). For detailed numerical results of all tested classifiers see Supplementary Fig. S3. Throughout this section we report each effect in the IKEM cohort alongside its counterpart under external validation (Section 2.5), where the classifiers trained and tuned on the IKEM dataset were applied to the independent NUDZ dataset. We quantify the transfer itself at the end of the section.

**Figure 1:**
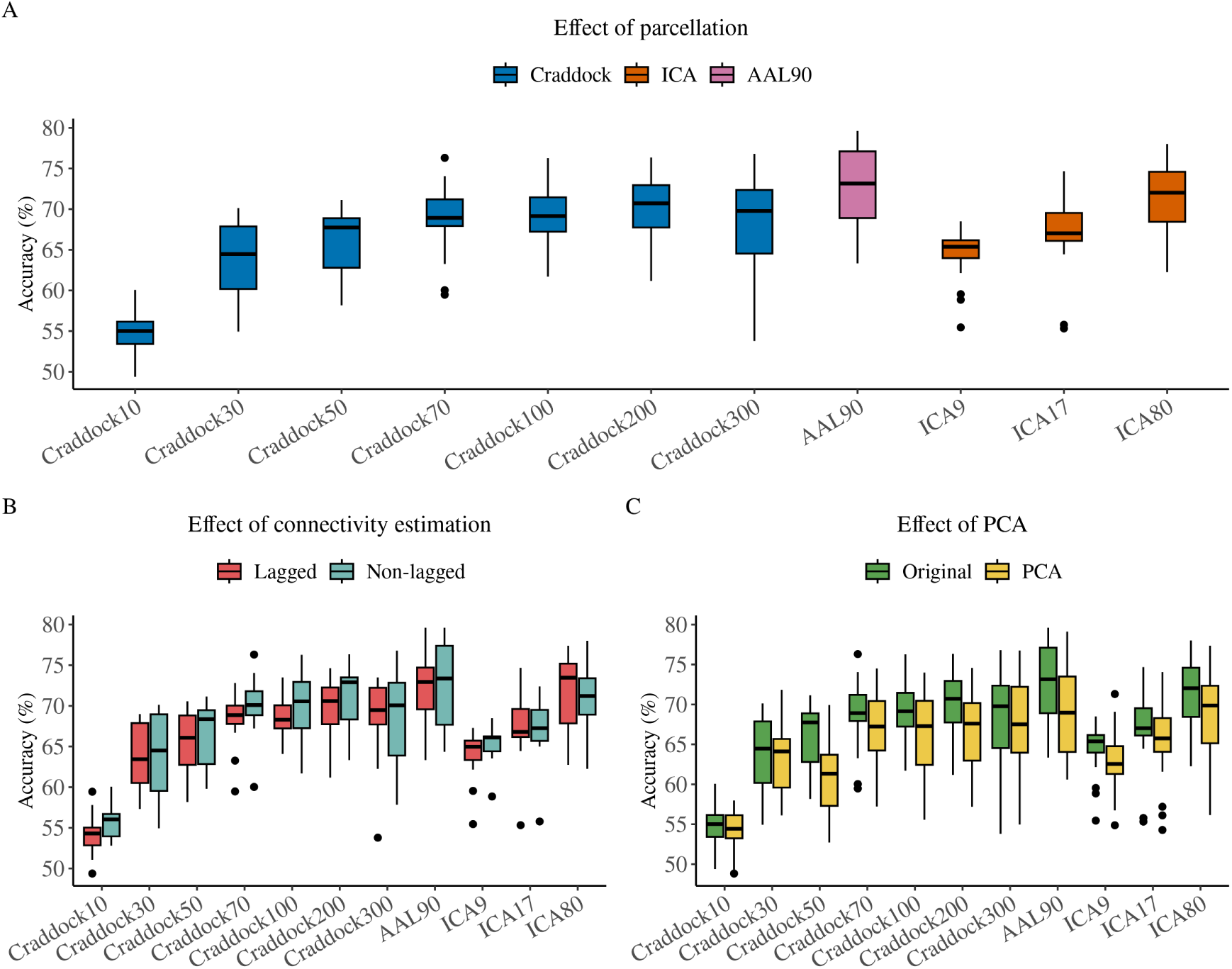
Accuracy across classifiers, per parcellation, connectivity type and PCA decomposition. A) Accuracy per parcellation, without PCA. B) Lagged against non-lagged connectivity, without PCA. C) With against without PCA. Each box in A and C summarises 34 values, the 17 tested classifiers on both connectivity types; each box in B summarises the 17 classifiers of one connectivity type.

**Table 1:** Performance of each parcellation, averaged over the 17 classifiers and, where both were run, both connectivity types; in parentheses, the same means on the PCA-reduced features. Craddock400, 500 and 950 were run non-lagged only, and 500 and 950 were run only with PCA (—) as running at original dimension size is infeasible.

| Parcellation | Accuracy | Sensitivity | Specificity | AUC-ROC |
| --- | --- | --- | --- | --- |
| Craddock10 | 54.9 (54.4) | 50.6 (50.3) | 59.3 (58.5) | 0.555 (0.564) |
| Craddock30 | 63.9 (63.2) | 61.7 (57.4) | 66.0 (69.0) | 0.693 (0.690) |
| Craddock50 | 66.1 (61.0) | 67.8 (53.9) | 64.5 (68.2) | 0.725 (0.678) |
| Craddock70 | 69.2 (66.9) | 69.1 (60.1) | 69.4 (73.7) | 0.767 (0.739) |
| Craddock100 | 69.2 (66.6) | 69.6 (58.8) | 68.8 (74.4) | 0.755 (0.729) |
| Craddock200 | 70.4 (66.7) | 71.8 (60.5) | 68.9 (72.9) | 0.774 (0.746) |
| Craddock300 | 68.7 (67.5) | 69.9 (60.0) | 67.4 (75.1) | 0.764 (0.754) |
| Craddock400 | 70.2 (68.8) | 70.9 (62.8) | 69.5 (74.9) | 0.771 (0.759) |
| Craddock500 | — (69.5) | — (64.4) | — (74.6) | — (0.759) |
| Craddock950 | — (68.7) | — (66.6) | — (70.8) | — (0.756) |
| AAL90 | 72.4 (69.3) | 73.5 (65.4) | 71.3 (73.1) | 0.787 (0.769) |
| ICA9 | 64.7 (62.9) | 63.3 (61.9) | 66.1 (64.0) | 0.696 (0.681) |
| ICA17 | 67.4 (65.9) | 67.2 (61.9) | 67.7 (69.9) | 0.736 (0.715) |
| ICA80 | 71.4 (68.7) | 72.8 (61.5) | 70.0 (75.8) | 0.786 (0.766) |

We found that classification performance depended primarily on the granularity of the parcellation (Fig. 1A; Fig. 2). Across the Craddock granularity levels, accuracy rose steeply across the coarsest levels and then saturated. Paired Wilcoxon signed-rank tests matched on classifier and connectivity type (Fig. 3; Supplementary Table S3) revealed pronounced gains from 10 to 30 regions (+9.9 percentage points, *p* = 1.6 *×* 10*^−^*^9^), from 30 to 50 (+1.5, *p* = 0.013) and from 50 to 70 (+2.6, *p* = 6.7 *×* 10*^−^*^5^). Beyond 70 regions, no two Craddock levels differed significantly in the IKEM cohort (70 against 100, 200 and 300: *p* = 0.81, 0.076 and 0.93; 100 against 200 and 300: *p* = 0.11 and 0.89; 200 against 300: *p* = 0.12). Craddock400, 500 and 950 were run on non-lagged features only and are therefore absent from these comparisons, which match on connectivity type. The results on external validation dataset showed the same steep rise and the same saturation point (10 to 30: +6.8; 30 to 50: +4.7; both *p <* 10*^−^*^9^; 50 to 70 and 70 to 100 not significant). Above it, however, that cohort retained modest further gains: +1.7 from 70 to 200 (*p* = 0.0038), +2.0 from 70 to 300 (*p* = 9.4 *×* 10*^−^*^5^), +0.6 from 100 to 200 (*p* = 0.049) and +0.9 from 100 to 300 (*p* = 0.011), with 200 and 300 again indistinguishable (*p* = 0.12). Accuracy therefore saturated at roughly 70 to 100 regions. At most two percentage points were available beyond that point, in one of the two cohorts, and we observed no decline at the finest levels.

**Figure 2:**
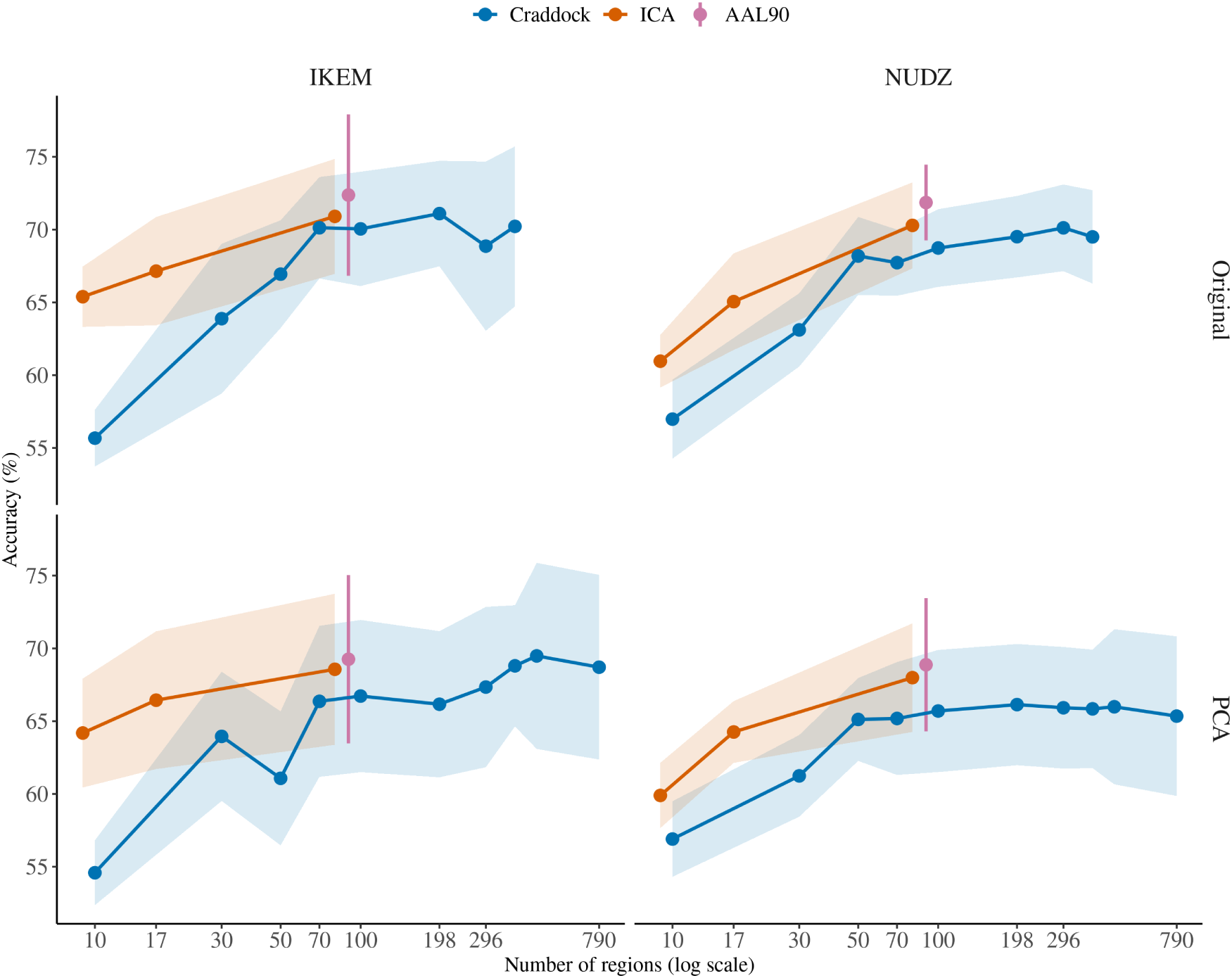
Accuracy against parcellation granularity, per cohort (columns) and feature space (rows). Points are means over the 17 tested classifiers, the ribbon is *±*1 SD; nonlagged connectivity only. The classifiers in original feature space stop at Craddock400, as running them is infeasible at the finest levels.

**Figure 3:**
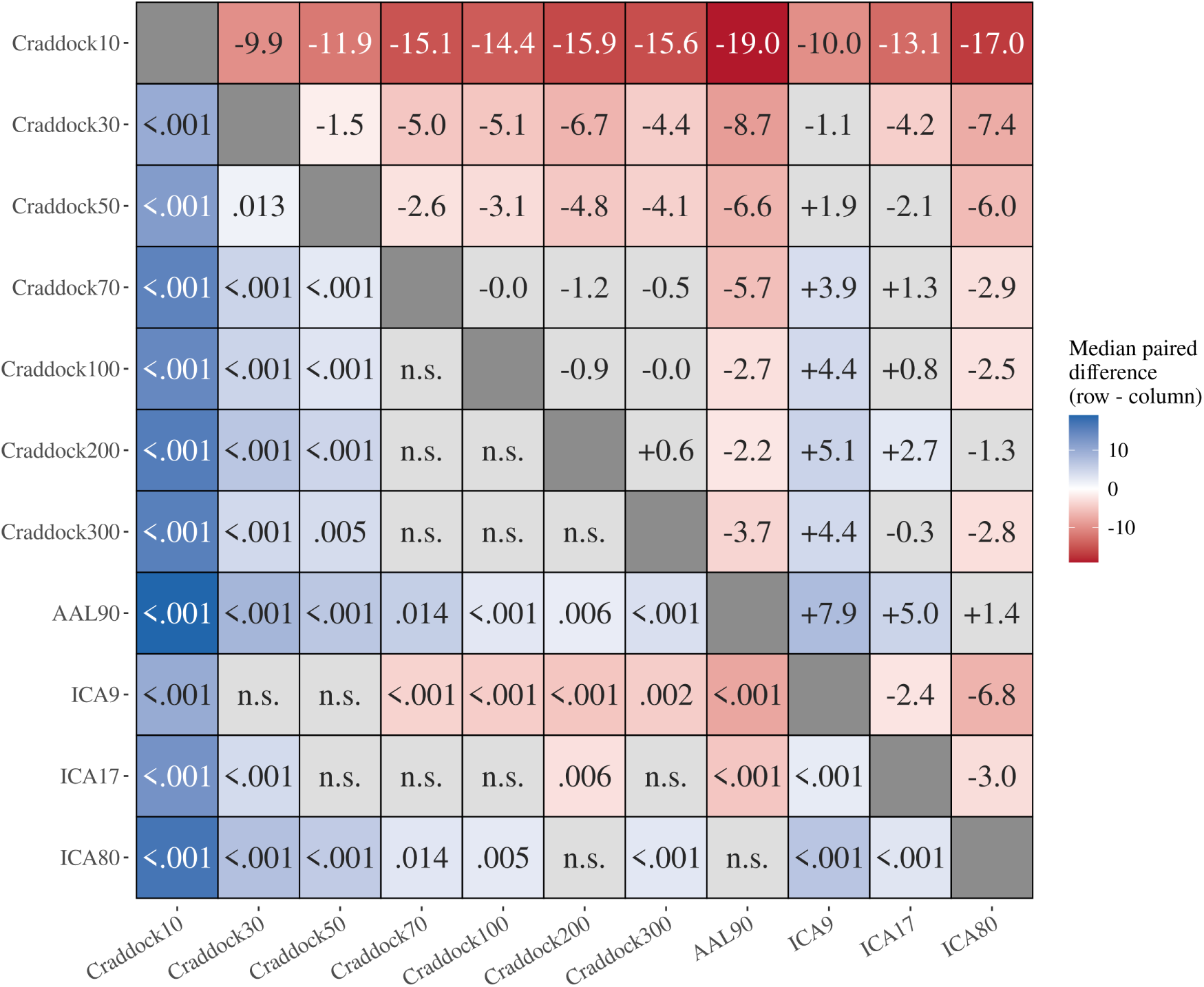
Pairwise comparisons between parcellations. Each cell is a paired Wilcoxon signed-rank test on the results without PCA, matched on classifier and connectivity type (FDR-adjusted). Colour encodes the median paired difference in accuracy between the row and the column parcellation: blue where the row performed better, red where it performed worse. The matrix is symmetric: cells above the diagonal give the median difference, cells below it the adjusted *p* for the same pair, written without its leading zero. Non-significant cells are grey and labelled n.s. in place of the *p*.

Above the plateau, the remaining differences between representations were small and did not follow the distinction between region-based and network-based features. AAL90 and ICA80, an anatomical atlas and a data-driven decomposition of comparable dimensionality, emerged as the two strongest representations. They did not differ significantly in the IKEM cohort (+1.4, *p* = 0.157), and AAL90 showed only a modest advantage under external validation (+1.6, *p* = 1.1 *×* 10*^−^*^4^). Their margin over the Craddock granularity levels was likewise modest, though consistent. AAL90, comprising 90 regions, outperformed every Craddock level tested without PCA, including levels several times finer: by +5.7 against Craddock70 (*p* = 0.014), +2.7 against Craddock100 (*p* = 2.3 *×* 10*^−^*^4^), +2.2 against Craddock200 (*p* = 0.0062) and +3.7 against Craddock300 (*p* = 7.9 *×* 10*^−^*^6^). The ordering was comparable in the independent cohort (+3.7, +3.3, +2.2 and +2.1; all *p <* 10*^−^*^5^). The ICA decompositions also sat above the Craddock curve at comparable dimensionality. ICA9 exceeded Craddock10 by 10.0 points and was indistinguishable from Craddock30 (*p* = 0.48), ICA17 exceeded Craddock30 by 4.2 (*p* = 5.4 *×* 10*^−^*^4^), and ICA80 exceeded Craddock70 and Craddock100 by 2.9 and 2.5 points (*p* = 0.014 and 0.0045) and Craddock300 by 2.8 (*p* = 5.4 *×* 10*^−^*^4^). Within the ICA representations, accuracy increased systematically with number of components, ICA9 yielding the weakest and ICA80 the strongest performance in both cohorts (all pairwise *p <* 10*^−^*^3^). The three decompositions thus fell on the same rising-then-saturating curve as the atlas-based representations. The single highest accuracy was obtained with an AAL90-based classifier (Supplementary Table S1).

We evaluated the two finest granularity levels, Craddock500 and Craddock950, on the PCA-reduced features alone, where the number of features is capped at 100 irrespective of the number of regions. No pair among Craddock300, 400, 500 and 950 differed significantly in either cohort (IKEM *p* = 0.28, 0.70 and 0.67 for successive levels; independent cohort *p* = 0.93, 0.62 and 0.091), and Craddock950 was indistinguishable from AAL90 in the IKEM cohort (*p* = 0.45; Supplementary Tables S9 and S10). However, a few comparisons within the plateau were significant, but the two cohorts pointed in different directions. In the IKEM cohort the finest levels won: Craddock400, 500 and 950 each beat Craddock70, 100 and 200 by 1.2 to 4.0 percentage points, in 8 of those 9 comparisons. In the NUDZ cohort AAL90 won instead, beating Craddock300, 400, 500 and 950 by 2.5, 2.2, 1.1 and 2.9 points (all *p <* 0.003). Extending granularity to very fine parcellations therefore neither recovered further accuracy nor degraded it substantially.

Next, we tested the effect of using lagged and non-lagged correlation to compute FC (Fig. 1B). Neither connectivity estimate systematically outperformed the other: no parcellation exhibited a significant difference in the IKEM cohort (smallest *p* = 0.10; Supplementary Table S5), and under external validation the only significant difference is at Craddock70, favouring lagged connectivity by 1.1 percentage points (*p* = 0.0042; Supplementary Table S6).

We found that PCA reduced accuracy significantly in 7 of the 11 parcellations in the IKEM cohort, and in 10 of the 11 under external validation, the sole exception in the latter being Craddock10 (Supplementary Tables S7 and S8; Fig. 1C). The same pattern held for AUC-ROC, with significant reductions in 6 and 10 parcellations respectively. In the IKEM cohort, AAL90 and ICA80 fell short of significance on AUC-ROC (*p* = 0.070 and 0.134), while ICA17 reached it on AUC-ROC but not on accuracy. Original features were therefore favoured on both metrics. Because the tuned number of components was capped at 100, the PCA-reduced features present the classifier with at most 100 features at every granularity level; the two settings are consequently comparable within themselves rather than against each other. Under PCA the IKEM granularity curve also exhibited a local dip at Craddock50, which fell 2.9 points below Craddock30 (61.0 against 63.2, *p* = 0.018; Fig. 2); the corresponding step was a gain of 1.5 points on the original features.

Finally, we quantified the cost of the transfer to the independent cohort (Table 2, Supplementary Table S2 and Fig. S4). It was modest: averaged across classifiers, AAL90 lost 0.6 and ICA80 lost 1.2 percentage points, while the finer Craddock levels changed by at most 1.2 points in either direction (Craddock300 rose from 68.7 internally to 69.9 under transfer, Craddock200 fell from 70.4 to 69.2). The best single AAL90-based model retained an accuracy of 76.1 (SD 2.7), against 79.6 (SD 8.2) internally, and the best Craddock300 model essentially held its value (76.0 against 76.8). Both the shape of the granularity curve and the ordering of the parcellations were thus reproduced at an independent site (Fig. 2; Fig. 4; Fig. 5).

**Figure 4:**
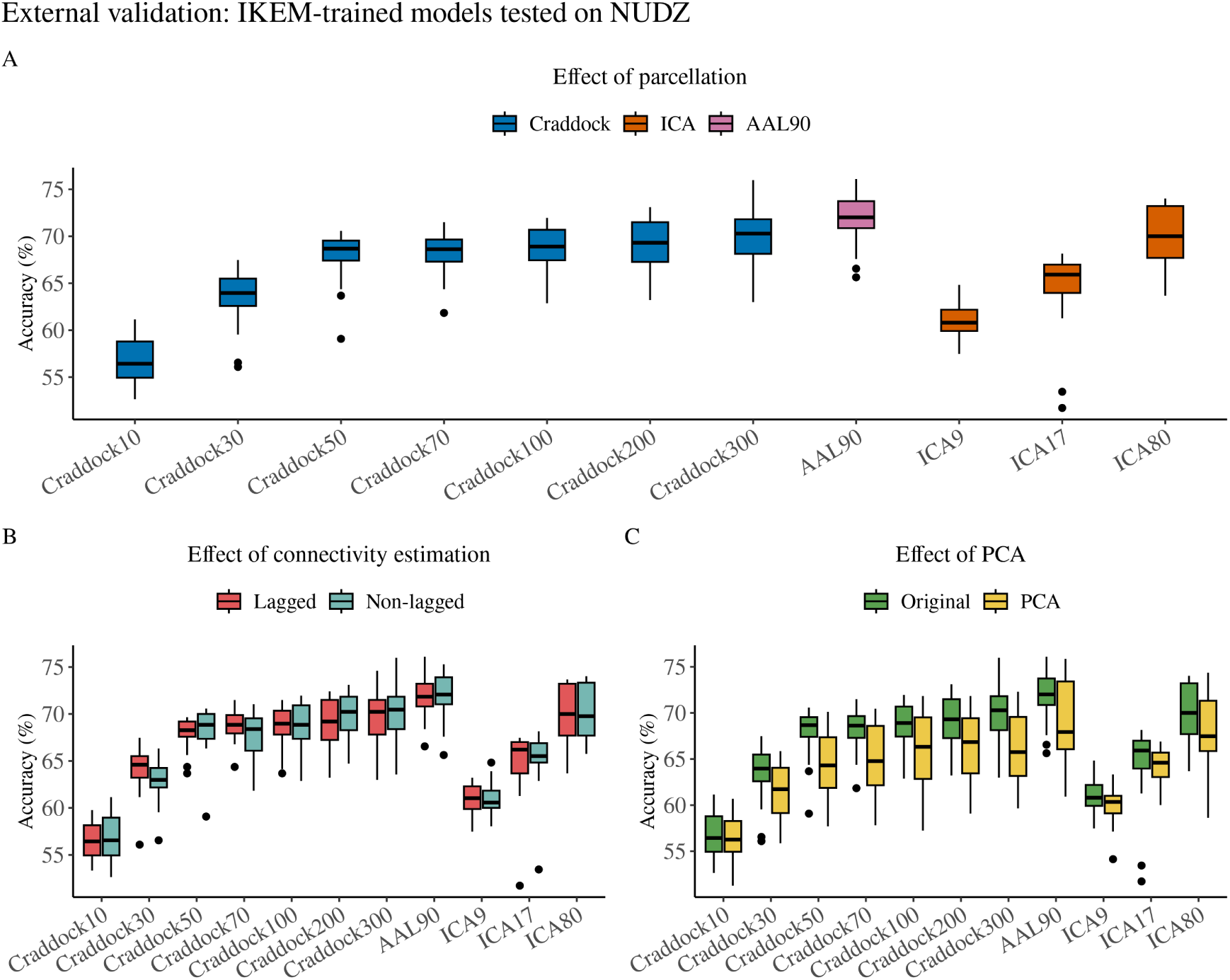
External validation (IKEM-trained models tested on NUDZ): accuracy across classifiers, per parcellation, connectivity type and PCA decomposition. A) Accuracy per parcellation, without PCA. B) Lagged against non-lagged connectivity, without PCA. C) With against without PCA. Each box in A and C summarises 34 values, the 17 tested classifiers on both connectivity types; each box in B summarises the 17 classifiers of one connectivity type.

**Figure 5:**
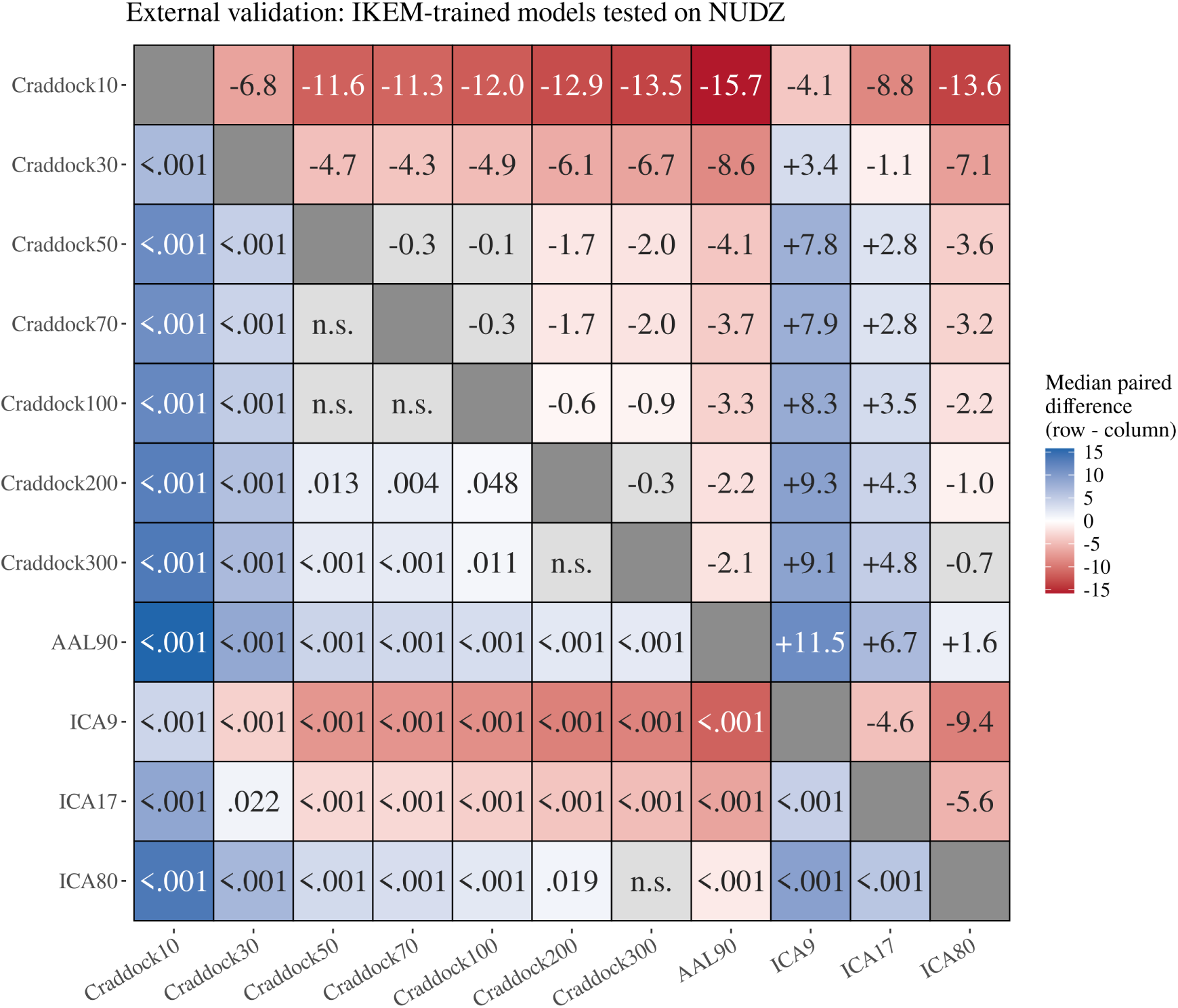
External validation (IKEM-trained models tested on NUDZ): pairwise comparisons between parcellations. Each cell is a paired Wilcoxon signed-rank test on the results without PCA, matched on classifier and connectivity type (FDR-adjusted). Colour encodes the median paired difference in accuracy between the row and the column parcellation: blue where the row performed better, red where it performed worse. The matrix is symmetric: cells above the diagonal give the median difference, cells below it the adjusted *p* for the same pair, written without its leading zero. Non-significant cells are grey and labelled n.s. in place of the *p*.

**Table 2:** External validation (train IKEM, test NUDZ): Performance of each parcellation, averaged over the 17 classifiers and, where both were run, both connectivity types; in parentheses, the same means on the PCA-reduced features. Craddock400, 500 and 950 were run non-lagged only, and 500 and 950 were run only with PCA (—) as running at original dimension size is infeasible.

| Parcellation | Accuracy | Sensitivity | Specificity | AUC-ROC |
| --- | --- | --- | --- | --- |
| Craddock10 | 56.7 (56.3) | 58.7 (59.5) | 54.6 (53.2) | 0.591 (0.589) |
| Craddock30 | 63.6 (61.4) | 66.6 (60.9) | 60.5 (61.9) | 0.693 (0.663) |
| Craddock50 | 68.1 (64.7) | 67.5 (56.5) | 68.6 (72.9) | 0.751 (0.713) |
| Craddock70 | 68.3 (65.3) | 69.8 (61.3) | 66.7 (69.2) | 0.752 (0.716) |
| Craddock100 | 68.6 (66.0) | 74.2 (65.6) | 62.9 (66.5) | 0.757 (0.724) |
| Craddock200 | 69.2 (66.2) | 76.9 (65.6) | 61.5 (66.8) | 0.768 (0.731) |
| Craddock300 | 69.9 (66.1) | 78.0 (64.1) | 61.7 (68.0) | 0.779 (0.734) |
| Craddock400 | 69.5 (65.8) | 75.7 (66.1) | 63.3 (65.6) | 0.775 (0.730) |
| Craddock500 | — (66.0) | — (67.0) | — (65.0) | — (0.732) |
| Craddock950 | — (65.3) | — (66.9) | — (63.8) | — (0.719) |
| AAL90 | 71.8 (69.2) | 77.4 (66.0) | 66.3 (72.4) | 0.807 (0.771) |
| ICA9 | 60.9 (59.9) | 75.6 (74.4) | 46.3 (45.5) | 0.672 (0.654) |
| ICA17 | 64.9 (64.2) | 72.1 (69.1) | 57.6 (59.3) | 0.716 (0.708) |
| ICA80 | 70.2 (68.0) | 75.4 (64.4) | 65.1 (71.7) | 0.772 (0.759) |

## 4 Discussion

### 4.1 Performance across parcellation strategies

Classification performance was largely driven by the granularity of the parcellation, not by the distinction between region-based and network-based features (Fig. 2; Fig. 3). Accuracy rose steeply from the coarsest levels and saturated at roughly 70 to 100 regions, after which further increase in granularity, up to Craddock950, brought no further gain. Above that resolution, how the regions are obtained matters little. An anatomical atlas and a data-driven ICA decomposition of similar dimensionality performed equally well in the IKEM cohort and differed by only 1.6 percentage points under external validation. The differences that remained between representations were a few percentage points, against the 11 to 15 points that came from granularity. The same pattern held in the NUDZ cohort (Fig. 5), so it is unlikely to be specific to a single site. At comparable granularity, region-based functional connectivity performed similarly as ICA-based functional network connectivity in terms of classification performance, so we see no evidence that the choice between them matters as much as granularity does.

ICA80 was included as a dimensionality-matched control, with its number of components chosen to approximate the dimensionality of the ROI atlases. That it matched the best atlas shows that, at similar dimensionality, the two representations carry similar discriminative information, and we found no sign that reducing the data to independent components discards information relevant for classification. The weaker performance of ICA9 and ICA17 is better explained by their coarseness than by their being network-based: accuracy rose with the number of components across the three ICA sets, and the Craddock granularity levels showed the same dependence within a single fixed scheme.

The three ICA sets differ, however, in more than the number of components, which limits what the comparison can show. ICA9 and ICA17 are hand-picked sets of resting-state networks. Components judged to be artefacts were discarded by visual inspection, and only the interpretable ones kept. This is a common practice in the functional network connectivity literature [Jafri et al., 2008, Arbabshirani et al., 2013, Damaraju et al., 2014]. In contrast, ICA80 is the raw output of the decomposition, kept in full, so it also includes components that such a selection would have thrown out and that would not usually be called networks at all. That the unselected set beat both curated ones suggests that components discarded as uninterpretable still carry information relevant to classification, or at least that hand-selection removes discriminative signal along with noise. ICA was therefore competitive only where it had given up one of its main advantages, i.e. a small set of recognisable networks.

Roughly 70 to 100 regions is thus enough to capture the connectivity differences between first-episode patients and controls at these sample sizes. Once granularity was matched, the scheme still mattered a little. Both AAL90, an anatomically defined atlas, and ICA80, a data-driven decomposition, ran a few percentage points above the Craddock granularity levels [Craddock et al., 2012] of similar size, and the ICA sets beat the corresponding Craddock levels throughout the coarse range. These margins are small beside the granularity effect, but they held in both cohorts, so something about how the regions themselves are created does matter. The schemes differ in several ways at once: how the boundaries are placed, how similar the regions are in size, and whether the regions are defined by anatomy or by the data. With one atlas and one decomposition method, we cannot separate these. Deciding between them would require parcellation method that varies one such property at a time.

However, despite comparable classification performance at comparable granularity, the two approaches also differ in what they cost to use. An ROI atlas is a fixed parcellation: the same regions apply to every participant, fold and site, at negligible computational cost. An ICA basis has to be estimated from the data and, to avoid leakage, re-estimated within every training fold, so the most expensive step of the pipeline is repeated once per fold. Its components also carry no fixed identity across decompositions, so relating a component to one derived from other data requires an explicit matching step [Arbabshirani et al., 2013]. At no or little gain in accuracy, an ROI parcellation might therefore be the more sensible default.

Finally, although this result may be anecdotal at the current stage and calls for a further validation across datasets and classification tasks, AAL90 was the best-performing representation overall, and the highest single accuracy was likewise obtained with it (Supplementary Table S1). It is a standard anatomical atlas [Tzourio-Mazoyer et al., 2002] and with 90 regions it sits right where accuracy plateaus, making it a reasonable default for this classification task. Functional parcellations of comparable granularity performed similarly and remain preferable where the functional interpretation of the regions matters.

### 4.2 Impact of additional design choices

We used Pearson correlation as our measure of functional connectivity, as it offers a simple and widely accepted approach for estimating statistical dependencies between time series. Prior work suggests that correlation is generally sufficient for capturing FC patterns relevant for clinical classification [Hlinka et al., 2011]. However, we acknowledge that more sophisticated transformations of the connectivity matrix have been proposed. Notably, the tangent space parametrisation of covariance matrices has shown marginally improved performance over correlation-based methods in benchmark studies such as that by Dadi et al. [2019].

Allowing the lag in the calculation of connectivity, as suggested by Jafri et al. [2008], did not affect classifier performance. We found non-lagged connectivity comparable to the lagged variants at every granularity level (Supplementary Table S5; Fig. 1B), and the single exception under external validation, a 1.1-point advantage for lagged connectivity at Craddock70, was isolated and not reproduced at any neighbouring level. Non-lagged functional connectivity is thus preferable in practice given its lower computational cost. Dimensionality reduction via PCA significantly reduced accuracy in 7 of the 11 parcellations in the IKEM cohort and in 10 of the 11 under external validation (Supplementary Tables S7 and S8; Fig. 1C). This convergent picture across metrics and cohorts indicates that PCA discards information relevant to discrimination, and PCA is therefore not recommended as a default preprocessing step for FC-based classification in this context. PCA was nonetheless informative here. By capping the classifier’s input at most 100 components irrespective of the number of regions, it holds input dimensionality fixed across the granularity levels, and the plateau emerged under this constraint as clearly as without it. The saturation we observed is thus a property of the connectivity estimates themselves, not of the difficulty of fitting a classifier to large number of features.

In terms of classifier choice, linear and kernel-based classifiers such as Ridge and SVM proved generally the most efficient (Supplementary Figs. S3 and S4), consistent with previous benchmarks [Arbabshirani et al., 2013, Dadi et al., 2019, Rashid and Calhoun, 2020] and supporting their use in other studies [Bučková et al., 2023]. The classifier mattered less than the representation, however. Once every model was tuned, median accuracy spanned 9.0 points between the best and the worst classifier in IKEM and 6.0 points under external validation, against 19.2 and 15.6 points between feature sets. Choosing the parcellation is therefore the more consequential decision, and a well-tuned linear model is enough at these sample sizes.

### 4.3 Generalisability and limitations

The design keeps the performance estimates conservative. Nested cross-validation with hyperparameter optimisation [Varoquaux et al., 2017, Varoquaux, 2018] avoids the variance inflation that leave-one-out cross-validation produces on small samples, and fitting the ICA separately within each training fold removes the leakage that occurs when spatial maps are estimated on the whole dataset. Every comparison is also paired on the classifier, so a difference between two parcellations has to hold across all 17 models rather than in a single favourable fit, and p-values were adjusted for multiple comparisons within each family of tests.

The real test of generalisability is the transfer to an independent dataset (Section 2.5). Transferring the models trained on the IKEM cohort to the NUDZ cohort cost little accuracy, and both the position of the saturation point and the ordering of the parcellations were preserved. AAL90 again ranked highest, and in that cohort it also beat ICA80, though by a small margin. The one concern is the shape of the plateau: where the IKEM cohort was flat above 70 regions, the NUDZ cohort kept small further gains of up to two percentage points at Craddock200 and Craddock300. The saturation point is therefore better described as a range of roughly 70 to 100 regions than as a single value. That the same pattern emerged at an independent site supports the generalisability of our findings. Validation on further independent datasets would permit a broader assessment of cross-site robustness.

### 4.4 Clinical relevance and future directions

Diagnosis is determined based on clinical interviews, but an independent validation method is not available. Using the resting-state functional connectivity has proven to be useful for the classification of healthy controls and patients with schizophrenia. Our findings further demonstrate that meaningful classification is indeed achievable in first-episode schizophrenia, a group often considered more difficult to classify due to subtler neural alterations [Wang et al., 2018a, Steardo et al., 2020].

Our results align well with prior literature on the classification of FES patients. While several studies have reported accuracies of 89.9% and higher [Wang et al., 2018a,b], these typically rely on small datasets, which limits their generalisability. In contrast, our study used a moderate sample size with rigorous nested cross-validation and hyperparameter optimisation. Our accuracies also fall below those reported for chronic patients [Kim et al., 2016, Cetin et al., 2016, Arbabshirani et al., 2013, 2014], which is expected: network-level disruptions are likely more pronounced later in the illness, whereas the differences we had to detect are those present at the first episode.

Compared to the task-based fMRI, the resting-state data is easier to obtain as it requires little to no effort from the participant and offers better comparability of the obtained results between studies [Skåtun et al., 2017, Orban et al., 2018]. In this respect, our results are encouraging and may be of practical value for early diagnosis, which is known to be crucial for optimal prognosis and treatment [Deco and Kringelbach, 2014]. Finally, while our study focused on functional connectivity, future work could explore multimodal approaches that integrate other neuroimaging modalities (structural MRI, DTI or EEG). Such multimodal classifiers could further boost performance and offer a more comprehensive characterisation of the brain changes induced by schizophrenia.

## 5 Conclusions

We compared functional connectivity features for the classification of first-episode schizophrenia. What mattered was the granularity of the parcellation, not whether its units were regions or networks. Accuracy rose steeply up to roughly 70 to 100 regions and then saturated, and a further order of magnitude, up to Craddock950, added nothing. Differences between representations of comparable size amounted to a few percentage points, against the 11 to 15 points attributable to granularity. Our attempt to replicate Arbabshirani et al. [2013] thus failed for a reason unrelated to ICA itself: the decomposition they used, reproduced here as ICA9, is too coarse and sat near the bottom of the curve in both cohorts. Only the finest and least curated decomposition, ICA80, matched the best atlas, while the coarse canonical variants, the ones the functional network connectivity literature actually interprets, performed worst. With no reliable accuracy advantage to set against the cost of estimating a basis within every fold and matching components across decompositions, we recommend region-based connectivity from a medium-resolution atlas such as AAL90, the best-performing representation in both cohorts. Whether this holds in larger independent datasets, and whether other imaging modalities add to it, remains to be tested.

## Author contributions

DT, DK, ZP, JHo, and JHl were responsible for conceptualization. DT, JHo, and JHl were responsible for data curation. DT, DK, ZP, and JHl were responsible for formal analysis. JHo, FS, and JHl were responsible for funding acquisition. DT, DK, ZP, and JHl were responsible for investigation. DT, DK, ZP, JHo, JT, FS, and JHl were responsible for methodology. FS and JHl were responsible for project administration. FS and JHl were responsible for resources. DT, DK, and ZP were responsible for software. JHl was responsible for supervision. DT and DK were responsible for validation. DT, DK, and JHl were responsible for visualization. DT, DK, ZP, and JHl were responsible for writing – original draft. DT, DK, ZP, JHo, JT, FS, and JHl were responsible for writing – review and editing.

## Acknowledgements

The publication was supported by ERDF-Project Brain dynamics, No. CZ.02.01.01/00/22_008/0004643, ERDF-Project Brainscape No. CZ.02.01.01/00/23_020/0008560, Czech Health Research Council Project No. NU21-08-00432, and PPPLZ - the Czech Academy of Sciences project PPLZ L100302451. Computational resources were provided by the e-INFRA CZ project (ID:90254), supported by the Ministry of Education, Youth and Sports of the Czech Republic.

## Data Availability Statement

The data that support the findings of this study are available from the corresponding author upon reasonable request.

## Code Availability Statement

The complete analysis code is publicly available at https://github.com/dominikklepl/ICA-Vs-ROI-Schizophrenia-Classification.

---

**Table S1:** Best classifier for each parcellation, with and without PCA; mean (SD) across five outer folds. Each entry is the maximum over 34 configurations (17 classifiers x lagged and non-lagged connectivity), selected on the same test folds it is reported on, and is therefore optimistic; it is given because it is the quantity comparable to published single-model results, while Table 1 gives the unbiased across-classifier mean. Craddock400, 500 and 950 were run non-lagged only, so their entries are a maximum over 17 configurations; 500 and 950 were run only with PCA.

| Parcellation | Arm | Model | Correlation type | Accuracy | Sensitivity | Specificity | AUC-ROC |
| --- | --- | --- | --- | --- | --- | --- | --- |
| Craddock10 | Original | XGBoost | Non-lagged | 60.1 (7.3) | 55.9 (16.0) | 64.2 (9.3) | 0.614 (0.066) |
| Craddock30 | Original | MLP | Non-lagged | 70.1 (7.6) | 65.0 (12.3) | 75.2 (11.0) | 0.773 (0.078) |
| Craddock50 | Original | XGBoost | Non-lagged | 71.1 (9.8) | 72.9 (4.8) | 69.3 (15.5) | 0.779 (0.109) |
| Craddock70 | Original | GradBoost | Non-lagged | 76.3 (4.8) | 75.2 (5.9) | 77.4 (9.8) | 0.823 (0.097) |
| Craddock100 | Original | ElasticNet | Non-lagged | 76.3 (9.2) | 79.7 (13.2) | 72.9 (7.3) | 0.802 (0.090) |
| Craddock200 | Original | SVM-Sigmoid | Non-lagged | 76.3 (8.7) | 77.5 (10.1) | 75.2 (11.6) | 0.829 (0.105) |
| Craddock300 | Original | Lasso | Non-lagged | 76.8 (8.8) | 76.3 (12.2) | 77.3 (12.3) | 0.845 (0.088) |
| Craddock400 | Original | SVM-Sigmoid | Non-lagged | 77.4 (8.2) | 78.5 (11.8) | 76.3 (9.4) | 0.844 (0.090) |
| AAL90 | Original | SVM-RBF | Lagged | 79.6 (8.2) | 81.8 (12.2) | 77.5 (11.3) | 0.840 (0.076) |
| ICA9 | Original | LightGBM | Non-lagged | 68.5 (5.6) | 66.2 (9.6) | 70.8 (2.2) | 0.708 (0.057) |
| ICA17 | Original | SVM-RBF | Lagged | 74.7 (4.2) | 71.8 (5.4) | 77.5 (5.0) | 0.790 (0.060) |
| ICA80 | Original | Lasso | Non-lagged | 78.0 (5.2) | 79.8 (11.4) | 76.2 (9.7) | 0.833 (0.064) |
| Craddock10 | PCA | XGBoost | Non-lagged | 58.0 (8.7) | 54.1 (15.2) | 61.9 (15.3) | 0.554 (0.054) |
| Craddock30 | PCA | MLP | Non-lagged | 71.8 (7.6) | 69.6 (11.6) | 74.1 (6.2) | 0.763 (0.079) |
| Craddock50 | PCA | SVM-Sigmoid | Non-lagged | 69.9 (13.6) | 70.5 (12.4) | 69.3 (15.9) | 0.740 (0.144) |
| Craddock70 | PCA | LDA | Lagged | 74.5 (11.7) | 72.8 (12.7) | 76.2 (12.0) | 0.804 (0.114) |
| Craddock100 | PCA | Ridge | Non-lagged | 74.0 (9.9) | 76.3 (9.7) | 71.6 (14.6) | 0.808 (0.140) |
| Craddock200 | PCA | SVM-Sigmoid | Non-lagged | 74.6 (7.9) | 71.7 (13.4) | 77.5 (9.5) | 0.819 (0.113) |
| Craddock300 | PCA | SVM-Linear | Lagged | 76.8 (12.1) | 72.9 (14.6) | 80.7 (12.2) | 0.813 (0.141) |
| Craddock400 | PCA | SVM-Sigmoid | Non-lagged | 74.5 (11.3) | 71.8 (12.6) | 77.3 (13.1) | 0.832 (0.113) |
| Craddock500 | PCA | Ridge | Non-lagged | 77.8 (12.9) | 77.3 (12.8) | 78.4 (17.4) | 0.826 (0.122) |
| Craddock950 | PCA | SVM-Linear | Non-lagged | 78.0 (8.9) | 78.4 (12.6) | 77.5 (12.7) | 0.832 (0.088) |
| AAL90 | PCA | SVM-Linear | Non-lagged | 79.1 (5.6) | 78.4 (10.4) | 79.8 (10.8) | 0.842 (0.063) |
| ICA9 | PCA | SVM-Poly | Non-lagged | 71.3 (5.6) | 53.9 (8.9) | 88.7 (5.2) | 0.755 (0.064) |
| ICA17 | PCA | SVM-RBF | Non-lagged | 74.1 (6.0) | 69.5 (9.9) | 78.6 (7.4) | 0.773 (0.084) |
| ICA80 | PCA | LDA | Lagged | 77.4 (8.1) | 78.5 (12.8) | 76.2 (12.0) | 0.836 (0.093) |

## Supplementary data

### Performance of every classifier and parcellation

The tables in this section give the numbers underlying the main-text figures. Table S1 and Table S2 report the single best classifier per parcellation in each cohort. These are maxima over 34 tasks selected on the same folds on which they are reported, and are therefore optimistic; they are given because they are the quantity comparable to the single-model accuracies usually published, whereas the unbiased summaries are the across-classifier means in Tables 1 and 2. The accuracy of every classifier at every parcellation is given in Figs. S3 and S4.

### Statistical comparisons

All comparisons are paired Wilcoxon signed-rank tests, matched on classifier, with p-values FDR-adjusted within each family of comparisons. Four families are reported per cohort: pairwise comparisons between parcellations on the original features (Tables S3 and S4, the statistics behind Figs. 3 and 5), lagged versus non-lagged connectivity (Tables S5 and S6), original versus PCA-reduced features (Tables S7 and S8), and pairwise comparisons between parcellations on the PCA-reduced features (Tables S9 and S10). The last family is separate because it is the only one that reaches the finest Craddock levels, which were run only with PCA.

**Table S2:** External validation (train IKEM, test NUDZ): Best classifier for each parcellation, with and without PCA; mean (SD) across the five IKEM-fold models. Each entry is the maximum over 34 configurations (17 classifiers x lagged and non-lagged connectivity), selected on the same test folds it is reported on, and is therefore optimistic; it is given because it is the quantity comparable to published single-model results, while Table 2 gives the unbiased across-classifier mean. Craddock400, 500 and 950 were run non-lagged only, so their entries are a maximum over 17 configurations; 500 and 950 were run only with PCA.

| Parcellation | Arm | Model | Correlation type | Accuracy | Sensitivity | Specificity | AUC-ROC |
| --- | --- | --- | --- | --- | --- | --- | --- |
| Craddock10 | Original | ElasticNet | Non-lagged | 61.1 (2.1) | 66.2 (10.6) | 56.1 (9.1) | 0.640 (0.012) |
| Craddock30 | Original | XGBoost | Lagged | 67.5 (1.7) | 71.7 (2.8) | 63.2 (5.4) | 0.734 (0.011) |
| Craddock50 | Original | AdaBoost | Non-lagged | 70.6 (1.6) | 72.9 (4.2) | 68.3 (4.5) | 0.767 (0.025) |
| Craddock70 | Original | LDA | Lagged | 71.5 (2.1) | 72.4 (5.2) | 70.6 (2.6) | 0.776 (0.030) |
| Craddock100 | Original | LDA | Non-lagged | 72.0 (2.9) | 77.9 (4.5) | 66.0 (5.0) | 0.780 (0.018) |
| Craddock200 | Original | Lasso | Non-lagged | 73.1 (2.3) | 84.8 (6.6) | 61.4 (4.6) | 0.825 (0.020) |
| Craddock300 | Original | Lasso | Non-lagged | 76.0 (0.9) | 86.9 (4.8) | 65.1 (4.5) | 0.848 (0.029) |
| Craddock400 | Original | Lasso | Non-lagged | 74.9 (2.4) | 85.7 (7.3) | 64.1 (5.4) | 0.845 (0.028) |
| AAL90 | Original | ElasticNet | Lagged | 76.1 (2.7) | 80.2 (8.8) | 72.0 (3.9) | 0.847 (0.040) |
| ICA9 | Original | KNN | Non-lagged | 64.8 (2.4) | 75.9 (2.1) | 53.8 (5.6) | 0.694 (0.026) |
| ICA17 | Original | ExtraTrees | Non-lagged | 68.2 (3.2) | 73.3 (2.5) | 63.0 (4.1) | 0.750 (0.016) |
| ICA80 | Original | SVM-Linear | Non-lagged | 74.0 (2.5) | 76.8 (6.3) | 71.3 (6.8) | 0.810 (0.015) |
| Craddock10 | PCA | Lasso | Non-lagged | 60.7 (2.6) | 65.1 (5.9) | 56.3 (7.6) | 0.636 (0.013) |
| Craddock30 | PCA | Ridge | Non-lagged | 65.9 (1.6) | 66.9 (4.9) | 64.8 (7.0) | 0.709 (0.016) |
| Craddock50 | PCA | LDA | Lagged | 70.1 (2.5) | 66.9 (7.5) | 73.3 (5.0) | 0.773 (0.027) |
| Craddock70 | PCA | Ridge | Non-lagged | 70.5 (2.6) | 69.7 (6.2) | 71.3 (5.3) | 0.781 (0.022) |
| Craddock100 | PCA | LDA | Lagged | 71.8 (4.0) | 78.6 (5.8) | 65.1 (4.2) | 0.789 (0.025) |
| Craddock200 | PCA | LDA | Lagged | 71.8 (2.2) | 77.0 (4.8) | 66.7 (5.3) | 0.796 (0.016) |
| Craddock300 | PCA | LDA | Lagged | 72.3 (3.0) | 82.1 (5.2) | 62.5 (4.3) | 0.822 (0.013) |
| Craddock400 | PCA | LDA | Non-lagged | 71.1 (2.9) | 77.0 (8.3) | 65.3 (7.5) | 0.808 (0.019) |
| Craddock500 | PCA | Ridge | Non-lagged | 73.3 (1.1) | 83.9 (5.0) | 62.8 (6.8) | 0.819 (0.021) |
| Craddock950 | PCA | Ridge | Non-lagged | 72.6 (2.1) | 81.6 (3.6) | 63.7 (4.9) | 0.807 (0.016) |
| AAL90 | PCA | ElasticNet | Non-lagged | 75.9 (2.2) | 80.7 (10.0) | 71.0 (8.3) | 0.833 (0.022) |
| ICA9 | PCA | SVM-Sigmoid | Non-lagged | 63.3 (2.2) | 77.5 (3.7) | 49.2 (4.5) | 0.698 (0.032) |
| ICA17 | PCA | SVM-Sigmoid | Lagged | 66.9 (3.8) | 70.3 (4.4) | 63.4 (4.4) | 0.734 (0.019) |
| ICA80 | PCA | Ridge | Non-lagged | 74.4 (3.4) | 76.8 (7.8) | 72.0 (6.7) | 0.810 (0.026) |

## Supplementary figures

**Table S3:** Pairwise Wilcoxon signed-rank tests of accuracy between parcellations, without PCA, paired on classifier and connectivity type (34 pairs per comparison). Median diff. is the median paired difference in percentage points, positive favouring Parcellation1. p-values FDR-adjusted across the 55 comparisons.

| Parcellation1 | Parcellation2 | W | Median diff. | p (FDR) | Parcellation1 | Parcellation2 | W | Median diff. | p (FDR) |
| --- | --- | --- | --- | --- | --- | --- | --- | --- | --- |
| Craddock10 | Craddock30 | 2.0 | -9.9 | 1.60e-09 | Craddock70 | Craddock200 | 186.5 | -1.2 | 7.55e-02 |
| Craddock10 | Craddock50 | 1.0 | -11.9 | 1.16e-09 | Craddock70 | Craddock300 | 291 | -0.5 | 9.26e-01 |
| Craddock10 | Craddock70 | 0.0 | -15.1 | 6.40e-10 | Craddock70 | AAL90 | 149 | -5.7 | 1.37e-02 |
| Craddock10 | Craddock100 | 0.0 | -14.4 | 6.40e-10 | Craddock70 | ICA9 | 556 | +3.9 | 2.02e-06 |
| Craddock10 | Craddock200 | 0.0 | -15.9 | 6.40e-10 | Craddock70 | ICA17 | 408 | +1.3 | 7.60e-02 |
| Craddock10 | Craddock300 | 0.0 | -15.6 | 6.40e-10 | Craddock70 | ICA80 | 150 | -2.9 | 1.41e-02 |
| Craddock10 | AAL90 | 0.0 | -19.0 | 6.40e-10 | Craddock100 | Craddock200 | 199.5 | -0.9 | 1.11e-01 |
| Craddock10 | ICA9 | 0.0 | -10.0 | 6.40e-10 | Craddock100 | Craddock300 | 307 | -0.0 | 8.85e-01 |
| Craddock10 | ICA17 | 0.0 | -13.1 | 6.40e-10 | Craddock100 | AAL90 | 85 | -2.7 | 2.32e-04 |
| Craddock10 | ICA80 | 0.0 | -17.0 | 6.40e-10 | Craddock100 | ICA9 | 559 | +4.4 | 1.38e-06 |
| Craddock30 | Craddock50 | 147.0 | -1.5 | 1.26e-02 | Craddock100 | ICA17 | 404 | +0.8 | 8.70e-02 |
| Craddock30 | Craddock70 | 30.0 | -5.0 | 5.84e-07 | Craddock100 | ICA80 | 127 | -2.5 | 4.52e-03 |
| Craddock30 | Craddock100 | 5.0 | -5.1 | 4.93e-09 | Craddock200 | Craddock300 | 394 | +0.6 | 1.15e-01 |
| Craddock30 | Craddock200 | 6.0 | -6.7 | 6.40e-09 | Craddock200 | AAL90 | 133 | -2.2 | 6.17e-03 |
| Craddock30 | Craddock300 | 16.0 | -4.4 | 6.14e-08 | Craddock200 | ICA9 | 580 | +5.1 | 5.44e-08 |
| Craddock30 | AAL90 | 0.0 | -8.7 | 6.40e-10 | Craddock200 | ICA17 | 461 | +2.7 | 6.17e-03 |
| Craddock30 | ICA9 | 252.5 | -1.1 | 4.75e-01 | Craddock200 | ICA80 | 192.5 | -1.3 | 8.93e-02 |
| Craddock30 | ICA17 | 96.0 | -4.2 | 5.41e-04 | Craddock300 | AAL90 | 50 | -3.7 | 7.93e-06 |
| Craddock30 | ICA80 | 0.0 | -7.4 | 6.40e-10 | Craddock300 | ICA9 | 482.5 | +4.4 | 1.71e-03 |
| Craddock50 | Craddock70 | 69.5 | -2.6 | 6.65e-05 | Craddock300 | ICA17 | 357 | -0.3 | 3.42e-01 |
| Craddock50 | Craddock100 | 55.0 | -3.1 | 1.33e-05 | Craddock300 | ICA80 | 95 | -2.8 | 5.37e-04 |
| Craddock50 | Craddock200 | 21.0 | -4.8 | 1.53e-07 | AAL90 | ICA9 | 571 | +7.9 | 2.57e-07 |
| Craddock50 | Craddock300 | 128.0 | -4.1 | 4.52e-03 | AAL90 | ICA17 | 526.5 | +5.0 | 5.40e-05 |
| Craddock50 | AAL90 | 29.0 | -6.6 | 5.30e-07 | AAL90 | ICA80 | 384.5 | +1.4 | 1.57e-01 |
| Craddock50 | ICA9 | 383.5 | +1.9 | 1.59e-01 | ICA9 | ICA17 | 100 | -2.4 | 7.04e-04 |
| Craddock50 | ICA17 | 198.0 | -2.1 | 1.09e-01 | ICA9 | ICA80 | 11 | -6.8 | 2.65e-08 |
| Craddock50 | ICA80 | 25.0 | -6.0 | 2.89e-07 | ICA17 | ICA80 | 74.5 | -3.0 | 9.02e-05 |
| Craddock70 | Craddock100 | 281.0 | -0.0 | 8.13e-01 |  |  |  |  |  |

**Table S4:** External validation (IKEM-trained models tested on NUDZ): Pairwise Wilcoxon signed-rank tests of accuracy between parcellations, without PCA, paired on classifier and connectivity type (34 pairs per comparison). Median diff. is the median paired difference in percentage points, positive favouring Parcellation1. p-values FDR-adjusted across the 55 comparisons.

| Parcellation1 | Parcellation2 | W | Median diff. | p (FDR) | Parcellation1 | Parcellation2 | W | Median diff. | p (FDR) |
| --- | --- | --- | --- | --- | --- | --- | --- | --- | --- |
| Craddock10 | Craddock30 | 1.0 | -6.8 | 6.40e-10 | Craddock70 | Craddock200 | 128.5 | -1.7 | 3.84e-03 |
| Craddock10 | Craddock50 | 0.0 | -11.6 | 4.00e-10 | Craddock70 | Craddock300 | 78 | -2.0 | 9.40e-05 |
| Craddock10 | Craddock70 | 0.0 | -11.3 | 4.00e-10 | Craddock70 | AAL90 | 0 | -3.7 | 4.00e-10 |
| Craddock10 | Craddock100 | 0.0 | -12.0 | 4.00e-10 | Craddock70 | ICA9 | 593 | +7.9 | 8.35e-10 |
| Craddock10 | Craddock200 | 0.0 | -12.9 | 4.00e-10 | Craddock70 | ICA17 | 539.5 | +2.8 | 1.07e-05 |
| Craddock10 | Craddock300 | 0.0 | -13.5 | 4.00e-10 | Craddock70 | ICA80 | 84 | -3.2 | 1.52e-04 |
| Craddock10 | AAL90 | 0.0 | -15.7 | 4.00e-10 | Craddock100 | Craddock200 | 180 | -0.6 | 4.85e-02 |
| Craddock10 | ICA9 | 3.0 | -4.1 | 1.23e-09 | Craddock100 | Craddock300 | 147 | -0.9 | 1.10e-02 |
| Craddock10 | ICA17 | 9.0 | -8.8 | 7.07e-09 | Craddock100 | AAL90 | 5 | -3.3 | 2.29e-09 |
| Craddock10 | ICA80 | 0.0 | -13.6 | 4.00e-10 | Craddock100 | ICA9 | 593 | +8.3 | 8.35e-10 |
| Craddock30 | Craddock50 | 0.0 | -4.7 | 4.00e-10 | Craddock100 | ICA17 | 532 | +3.5 | 2.37e-05 |
| Craddock30 | Craddock70 | 0.0 | -4.3 | 4.00e-10 | Craddock100 | ICA80 | 99.5 | -2.2 | 5.19e-04 |
| Craddock30 | Craddock100 | 1.0 | -4.9 | 6.40e-10 | Craddock200 | Craddock300 | 204 | -0.3 | 1.22e-01 |
| Craddock30 | Craddock200 | 1.0 | -6.1 | 6.40e-10 | Craddock200 | AAL90 | 2 | -2.2 | 8.35e-10 |
| Craddock30 | Craddock300 | 0.0 | -6.7 | 4.00e-10 | Craddock200 | ICA9 | 595 | +9.3 | 4.00e-10 |
| Craddock30 | AAL90 | 0.0 | -8.6 | 4.00e-10 | Craddock200 | ICA17 | 557 | +4.3 | 1.31e-06 |
| Craddock30 | ICA9 | 517.0 | +3.4 | 9.40e-05 | Craddock200 | ICA80 | 159 | -1.0 | 1.91e-02 |
| Craddock30 | ICA17 | 161.5 | -1.1 | 2.17e-02 | Craddock300 | AAL90 | 41 | -2.1 | 2.03e-06 |
| Craddock30 | ICA80 | 1.0 | -7.1 | 6.40e-10 | Craddock300 | ICA9 | 592 | +9.1 | 1.23e-09 |
| Craddock50 | Craddock70 | 248.5 | -0.3 | 4.14e-01 | Craddock300 | ICA17 | 571 | +4.8 | 1.57e-07 |
| Craddock50 | Craddock100 | 231.5 | -0.1 | 2.78e-01 | Craddock300 | ICA80 | 233 | -0.7 | 2.81e-01 |
| Craddock50 | Craddock200 | 151.0 | -1.7 | 1.30e-02 | AAL90 | ICA9 | 595 | +11.5 | 4.00e-10 |
| Craddock50 | Craddock300 | 76.0 | -2.0 | 8.16e-05 | AAL90 | ICA17 | 592 | +6.7 | 1.23e-09 |
| Craddock50 | AAL90 | 0.0 | -4.1 | 4.00e-10 | AAL90 | ICA80 | 514.5 | +1.6 | 1.14e-04 |
| Craddock50 | ICA9 | 585.0 | +7.8 | 9.18e-09 | ICA9 | ICA17 | 65 | -4.6 | 2.84e-05 |
| Craddock50 | ICA17 | 510.0 | +2.8 | 1.62e-04 | ICA9 | ICA80 | 0 | -9.4 | 4.00e-10 |
| Craddock50 | ICA80 | 102.0 | -3.6 | 6.16e-04 | ICA17 | ICA80 | 5 | -5.6 | 2.29e-09 |
| Craddock70 | Craddock100 | 211.5 | -0.3 | 1.60e-01 |  |  |  |  |  |

**Table S5:** Wilcoxon signed-rank tests of lagged against non-lagged connectivity, per parcellation, without PCA, paired on classifier (17 pairs). Median diff. is the median paired difference in accuracy, in percentage points. p-values FDR-adjusted.

| Space | FC type 1 | FC type 2 | W | Median diff. | p (FDR) |
| --- | --- | --- | --- | --- | --- |
| Craddock10 | Lagged | Non-lagged | 34.0 | -1.7 | 0.102 |
| Craddock30 | Lagged | Non-lagged | 74.0 | -0.5 | 0.927 |
| Craddock50 | Lagged | Non-lagged | 39.0 | -1.1 | 0.146 |
| Craddock70 | Lagged | Non-lagged | 24.5 | -1.2 | 0.102 |
| Craddock100 | Lagged | Non-lagged | 28.0 | -1.1 | 0.102 |
| Craddock200 | Lagged | Non-lagged | 33.0 | -1.1 | 0.102 |
| Craddock300 | Lagged | Non-lagged | 64.0 | -1.1 | 0.777 |
| AAL90 | Lagged | Non-lagged | 69.0 | -0.5 | 0.831 |
| ICA9 | Lagged | Non-lagged | 32.0 | -1.1 | 0.102 |
| ICA17 | Lagged | Non-lagged | 87.0 | +0.0 | 0.777 |
| ICA80 | Lagged | Non-lagged | 111.0 | +1.1 | 0.171 |

**Table S6:**
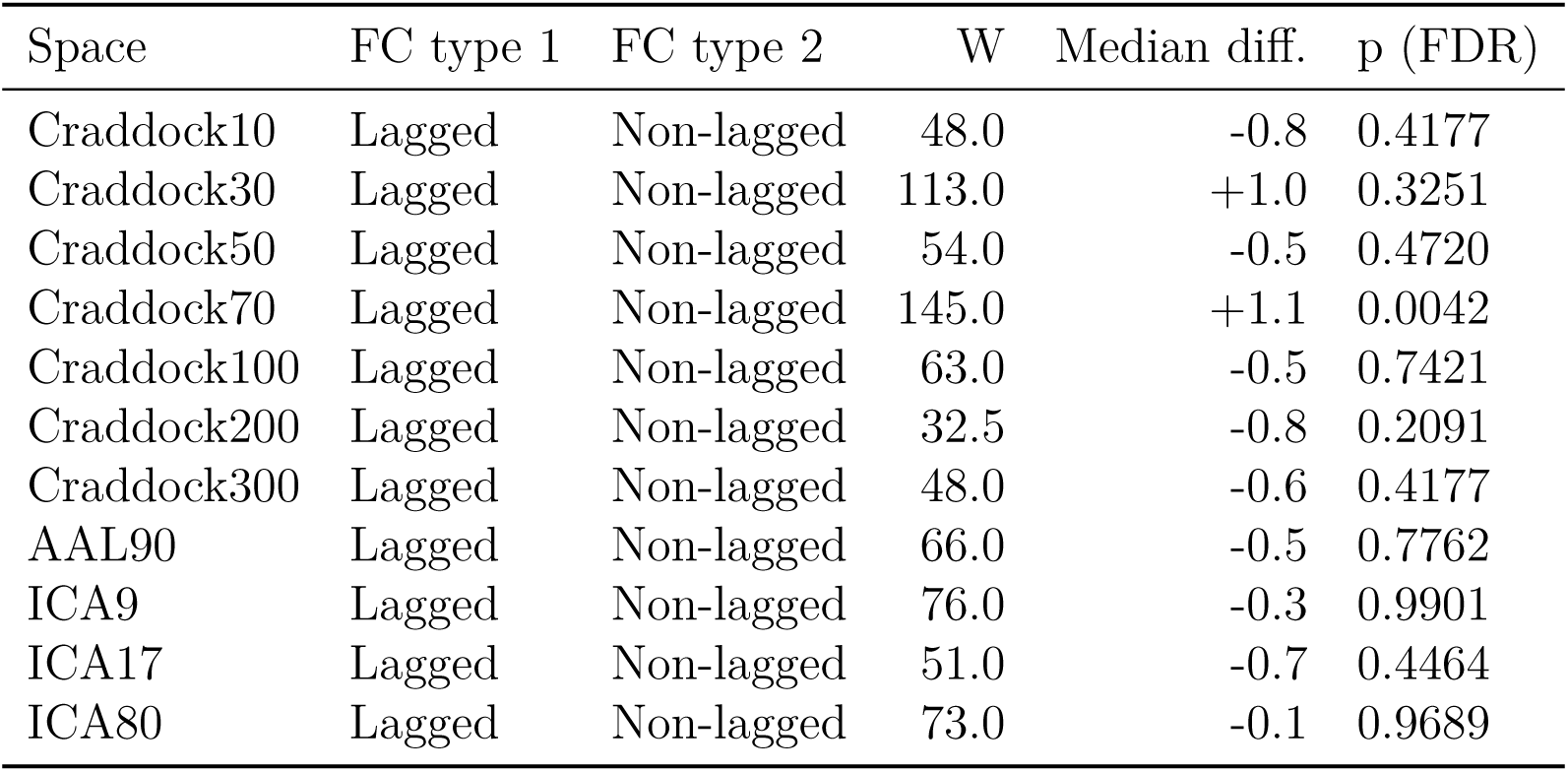
External validation (IKEM-trained models tested on NUDZ): Wilcoxon signed-rank tests of lagged against non-lagged connectivity, per parcellation, without PCA, paired on classifier (17 pairs). Median diff. is the median paired difference in accuracy, in percentage points. p-values FDR-adjusted.

| Space | FC type 1 | FC type 2 | W | Median diff. | p (FDR) |
| --- | --- | --- | --- | --- | --- |
| Craddock10 | Lagged | Non-lagged | 48.0 | -0.8 | 0.4177 |
| Craddock30 | Lagged | Non-lagged | 113.0 | +1.0 | 0.3251 |
| Craddock50 | Lagged | Non-lagged | 54.0 | -0.5 | 0.4720 |
| Craddock70 | Lagged | Non-lagged | 145.0 | +1.1 | 0.0042 |
| Craddock100 | Lagged | Non-lagged | 63.0 | -0.5 | 0.7421 |
| Craddock200 | Lagged | Non-lagged | 32.5 | -0.8 | 0.2091 |
| Craddock300 | Lagged | Non-lagged | 48.0 | -0.6 | 0.4177 |
| AAL90 | Lagged | Non-lagged | 66.0 | -0.5 | 0.7762 |
| ICA9 | Lagged | Non-lagged | 76.0 | -0.3 | 0.9901 |
| ICA17 | Lagged | Non-lagged | 51.0 | -0.7 | 0.4464 |
| ICA80 | Lagged | Non-lagged | 73.0 | -0.1 | 0.9689 |

**Table S7:** Wilcoxon signed-rank tests of original against PCA-reduced features, per parcellation, for accuracy and AUC-ROC. p-values FDR-adjusted within each metric.

| Space | W (BA) | p.adj (BA) | W (AUC) | p.adj (AUC) |
| --- | --- | --- | --- | --- |
| Craddock10 | 346.0 | 4.58e-01 | 227 | 0.286850 |
| Craddock30 | 328.0 | 6.09e-01 | 294 | 0.973081 |
| Craddock50 | 566.0 | 2.23e-06 | 529 | 0.000233 |
| Craddock70 | 426.0 | 4.24e-02 | 449 | 0.023620 |
| Craddock100 | 490.0 | 2.26e-03 | 437 | 0.029514 |
| Craddock200 | 529.0 | 1.14e-04 | 512 | 0.000573 |
| Craddock300 | 353.5 | 4.22e-01 | 300 | 0.973081 |
| AAL90 | 461.0 | 1.18e-02 | 415 | 0.069885 |
| ICA9 | 431.0 | 3.96e-02 | 438 | 0.029514 |
| ICA17 | 406.0 | 8.76e-02 | 489 | 0.002478 |
| ICA80 | 434.0 | 3.96e-02 | 395 | 0.134245 |

**Table S8:** External validation (IKEM-trained models tested on NUDZ): Wilcoxon signed-rank tests of original against PCA-reduced features, per parcellation, for accuracy and AUC-ROC. p-values FDR-adjusted within each metric.

| Space | W (BA) | p.adj (BA) | W (AUC) | p.adj (AUC) |
| --- | --- | --- | --- | --- |
| Craddock10 | 325.0 | 6.27e-01 | 298 | 1.00e+00 |
| Craddock30 | 529.5 | 4.26e-05 | 542 | 1.13e-05 |
| Craddock50 | 554.0 | 3.10e-06 | 540 | 1.19e-05 |
| Craddock70 | 573.0 | 2.19e-07 | 568 | 1.61e-06 |
| Craddock100 | 502.5 | 3.19e-04 | 537 | 1.24e-05 |
| Craddock200 | 581.0 | 1.34e-07 | 555 | 2.78e-06 |
| Craddock300 | 575.5 | 2.06e-07 | 560 | 1.84e-06 |
| AAL90 | 525.0 | 5.67e-05 | 563 | 1.77e-06 |
| ICA9 | 473.5 | 2.30e-03 | 538 | 1.24e-05 |
| ICA17 | 452.0 | 7.91e-03 | 504 | 2.57e-04 |
| ICA80 | 505.0 | 2.97e-04 | 449 | 9.45e-03 |

**Table S9:**
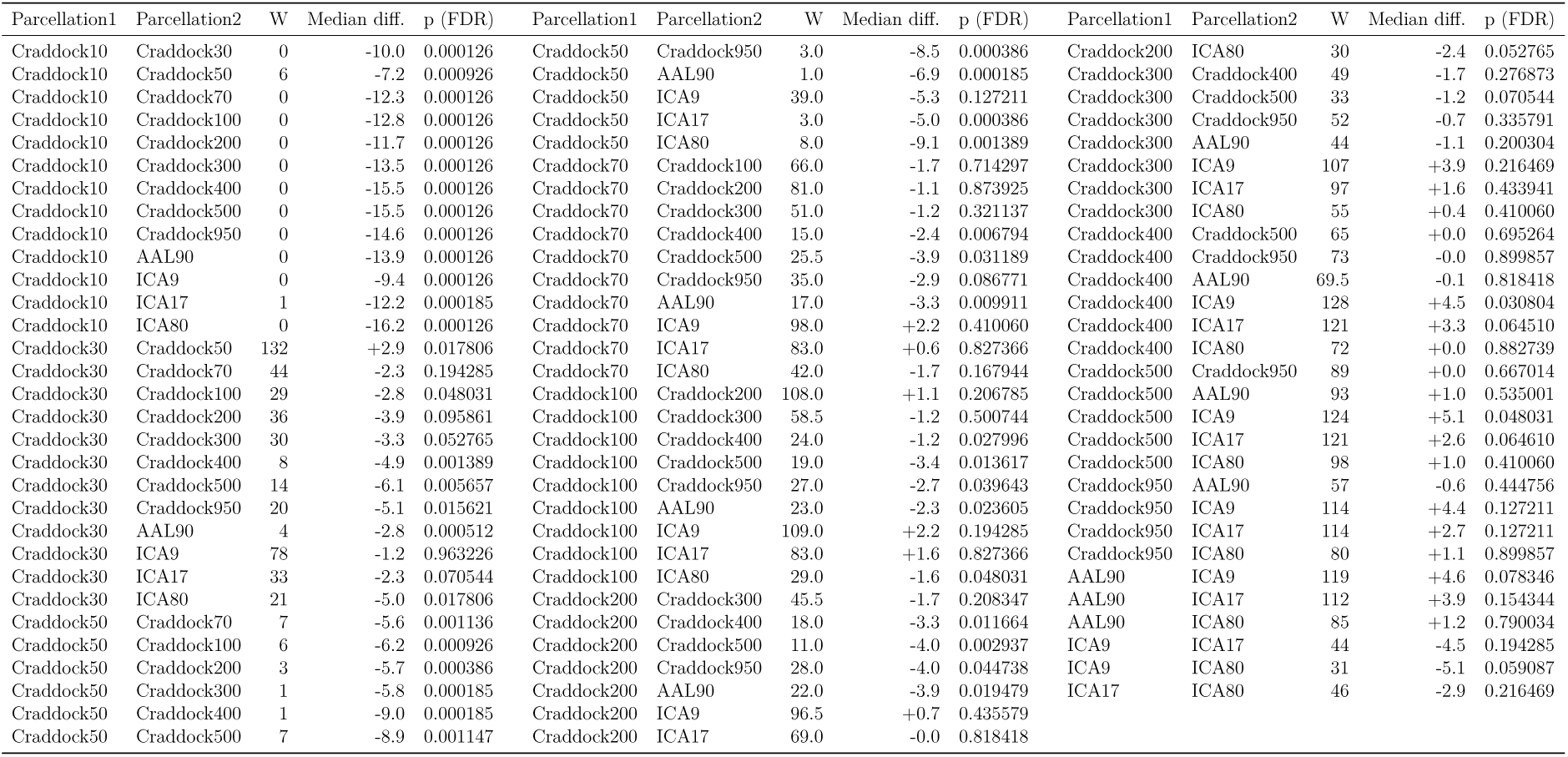
Pairwise Wilcoxon signed-rank tests of accuracy between parcellations on the PCA-reduced, non-lagged features, paired on classifier (17 pairs per comparison). PCA is the only setting run at every granularity level, so this is the comparison that covers all of them, Craddock500 and Craddock950 included. Median diff. as in Table S3. p-values FDR-adjusted within this family, separately from the tests without PCA.

**Table S10:** External validation (IKEM-trained models tested on NUDZ): Pairwise Wilcoxon signed-rank tests of accuracy between parcellations on the PCA-reduced, non-lagged features, paired on classifier (17 pairs per comparison). PCA is the only setting run at every granularity level, so this is the comparison that covers all of them, Craddock500 and Craddock950 included. Median diff. as in Table S4. p-values FDR-adjusted within this family, separately from the tests without PCA.

| Parcellation1 | Parcellation2 | W | Median diff. | p (FDR) | Parcellation1 | Parcellation2 | W | Median diff. | p (FDR) | Parcellation1 | Parcellation2 | W | Median diff. | p (FDR) |
| --- | --- | --- | --- | --- | --- | --- | --- | --- | --- | --- | --- | --- | --- | --- |
| Craddock10 | Craddock30 | 4.0 | -4.8 | 3.14e-04 | Craddock50 | Craddock950 | 65.5 | -0.6 | 6.79e-01 | Craddock200 | ICA80 | 35 | -2.4 | 8.42e-02 |
| Craddock10 | Craddock50 | 0.0 | -8.4 | 6.94e-05 | Craddock50 | AAL90 | 0.0 | -3.8 | 6.94e-05 | Craddock300 | Craddock400 | 79 | +0.1 | 9.28e-01 |
| Craddock10 | Craddock70 | 0.0 | -8.5 | 6.94e-05 | Craddock50 | ICA9 | 153.0 | +6.3 | 6.94e-05 | Craddock300 | Craddock500 | 73 | +0.2 | 9.01e-01 |
| Craddock10 | Craddock100 | 0.0 | -8.6 | 6.94e-05 | Craddock50 | ICA17 | 114.0 | +1.1 | 1.21e-01 | Craddock300 | Craddock950 | 92 | +0.5 | 5.70e-01 |
| Craddock10 | Craddock200 | 0.0 | -9.9 | 6.94e-05 | Craddock50 | ICA80 | 15.0 | -3.7 | 4.37e-03 | Craddock300 | AAL90 | 4 | -2.5 | 3.14e-04 |
| Craddock10 | Craddock300 | 0.0 | -9.5 | 6.94e-05 | Craddock70 | Craddock100 | 40.0 | -0.7 | 1.48e-01 | Craddock300 | ICA9 | 149 | +6.0 | 3.14e-04 |
| Craddock10 | Craddock400 | 0.0 | -9.1 | 6.94e-05 | Craddock70 | Craddock200 | 48.0 | -1.3 | 2.53e-01 | Craddock300 | ICA17 | 114 | +2.0 | 1.21e-01 |
| Craddock10 | Craddock500 | 0.0 | -9.3 | 6.94e-05 | Craddock70 | Craddock300 | 42.0 | -1.0 | 1.61e-01 | Craddock300 | ICA80 | 25 | -2.9 | 2.34e-02 |
| Craddock10 | Craddock950 | 0.0 | -9.8 | 6.94e-05 | Craddock70 | Craddock400 | 49.0 | -0.7 | 2.63e-01 | Craddock400 | Craddock500 | 63 | -1.0 | 6.24e-01 |
| Craddock10 | AAL90 | 0.0 | -12.9 | 6.94e-05 | Craddock70 | Craddock500 | 50.0 | -1.4 | 2.88e-01 | Craddock400 | Craddock950 | 84 | +0.3 | 7.62e-01 |
| Craddock10 | ICA9 | 7.0 | -3.0 | 6.94e-04 | Craddock70 | Craddock950 | 68.0 | -1.4 | 7.53e-01 | Craddock400 | AAL90 | 5.5 | -2.2 | 4.36e-04 |
| Craddock10 | ICA17 | 0.0 | -7.2 | 6.94e-05 | Craddock70 | AAL90 | 2.0 | -3.7 | 1.81e-04 | Craddock400 | ICA9 | 149 | +7.1 | 3.14e-04 |
| Craddock10 | ICA80 | 0.0 | -10.1 | 6.94e-05 | Craddock70 | ICA9 | 150.0 | +6.4 | 2.78e-04 | Craddock400 | ICA17 | 116 | +2.6 | 1.02e-01 |
| Craddock30 | Craddock50 | 0.0 | -3.6 | 6.94e-05 | Craddock70 | ICA17 | 104.5 | +1.3 | 2.60e-01 | Craddock400 | ICA80 | 27 | -2.3 | 3.10e-02 |
| Craddock30 | Craddock70 | 4.0 | -4.6 | 3.14e-04 | Craddock70 | ICA80 | 9.0 | -3.0 | 1.11e-03 | Craddock500 | Craddock950 | 117 | +0.7 | 9.09e-02 |
| Craddock30 | Craddock100 | 4.0 | -5.6 | 3.14e-04 | Craddock100 | Craddock200 | 61.0 | -0.1 | 5.70e-01 | Craddock500 | AAL90 | 13 | -1.1 | 2.56e-03 |
| Craddock30 | Craddock200 | 3.0 | -4.8 | 2.78e-04 | Craddock100 | Craddock300 | 61.5 | -0.1 | 5.70e-01 | Craddock500 | ICA9 | 145 | +8.2 | 7.83e-04 |
| Craddock30 | Craddock300 | 2.0 | -5.1 | 1.81e-04 | Craddock100 | Craddock400 | 58.0 | +0.0 | 5.70e-01 | Craddock500 | ICA17 | 111 | +0.8 | 1.48e-01 |
| Craddock30 | Craddock400 | 5.0 | -5.3 | 4.08e-04 | Craddock100 | Craddock500 | 61.5 | -0.5 | 5.70e-01 | Craddock500 | ICA80 | 24 | -1.0 | 2.04e-02 |
| Craddock30 | Craddock500 | 10.0 | -6.0 | 1.42e-03 | Craddock100 | Craddock950 | 81.5 | +0.0 | 8.46e-01 | Craddock950 | AAL90 | 0 | -2.9 | 6.94e-05 |
| Craddock30 | Craddock950 | 12.0 | -4.9 | 2.21e-03 | Craddock100 | AAL90 | 0.0 | -3.2 | 6.94e-05 | Craddock950 | ICA9 | 141 | +6.3 | 2.21e-03 |
| Craddock30 | AAL90 | 0.0 | -8.3 | 6.94e-05 | Craddock100 | ICA9 | 147.0 | +7.1 | 5.25e-04 | Craddock950 | ICA17 | 97 | +0.3 | 4.25e-01 |
| Craddock30 | ICA9 | 112.5 | +0.7 | 1.20e-01 | Craddock100 | ICA17 | 117.0 | +1.7 | 9.03e-02 | Craddock950 | ICA80 | 27 | -3.0 | 3.27e-02 |
| Craddock30 | ICA17 | 13.0 | -3.0 | 2.66e-03 | Craddock100 | ICA80 | 19.0 | -2.3 | 8.50e-03 | AAL90 | ICA9 | 153 | +9.5 | 6.94e-05 |
| Craddock30 | ICA80 | 1.0 | -7.2 | 1.32e-04 | Craddock200 | Craddock300 | 86.0 | +0.3 | 7.34e-01 | AAL90 | ICA17 | 147 | +4.1 | 5.25e-04 |
| Craddock50 | Craddock70 | 75.5 | -0.3 | 9.73e-01 | Craddock200 | Craddock400 | 87.5 | +0.3 | 6.79e-01 | AAL90 | ICA80 | 89.5 | -0.2 | 6.24e-01 |
| Craddock50 | Craddock100 | 46.0 | -0.6 | 2.30e-01 | Craddock200 | Craddock500 | 72.5 | -0.8 | 8.92e-01 | ICA9 | ICA17 | 0 | -4.9 | 6.94e-05 |
| Craddock50 | Craddock200 | 43.0 | -1.5 | 1.67e-01 | Craddock200 | Craddock950 | 93.0 | +0.2 | 5.55e-01 | ICA9 | ICA80 | 0 | -7.9 | 6.94e-05 |
| Craddock50 | Craddock300 | 49.0 | -0.1 | 2.63e-01 | Craddock200 | AAL90 | 5.0 | -2.8 | 4.08e-04 | ICA17 | ICA80 | 9 | -3.7 | 1.12e-03 |
| Craddock50 | Craddock400 | 43.0 | -0.7 | 1.72e-01 | Craddock200 | ICA9 | 148.0 | +7.2 | 4.08e-04 |  |  |  |  |  |
| Craddock50 | Craddock500 | 51.0 | -2.2 | 3.02e-01 | Craddock200 | ICA17 | 117.0 | +2.2 | 9.24e-02 |  |  |  |  |  |

**Figure S1:**
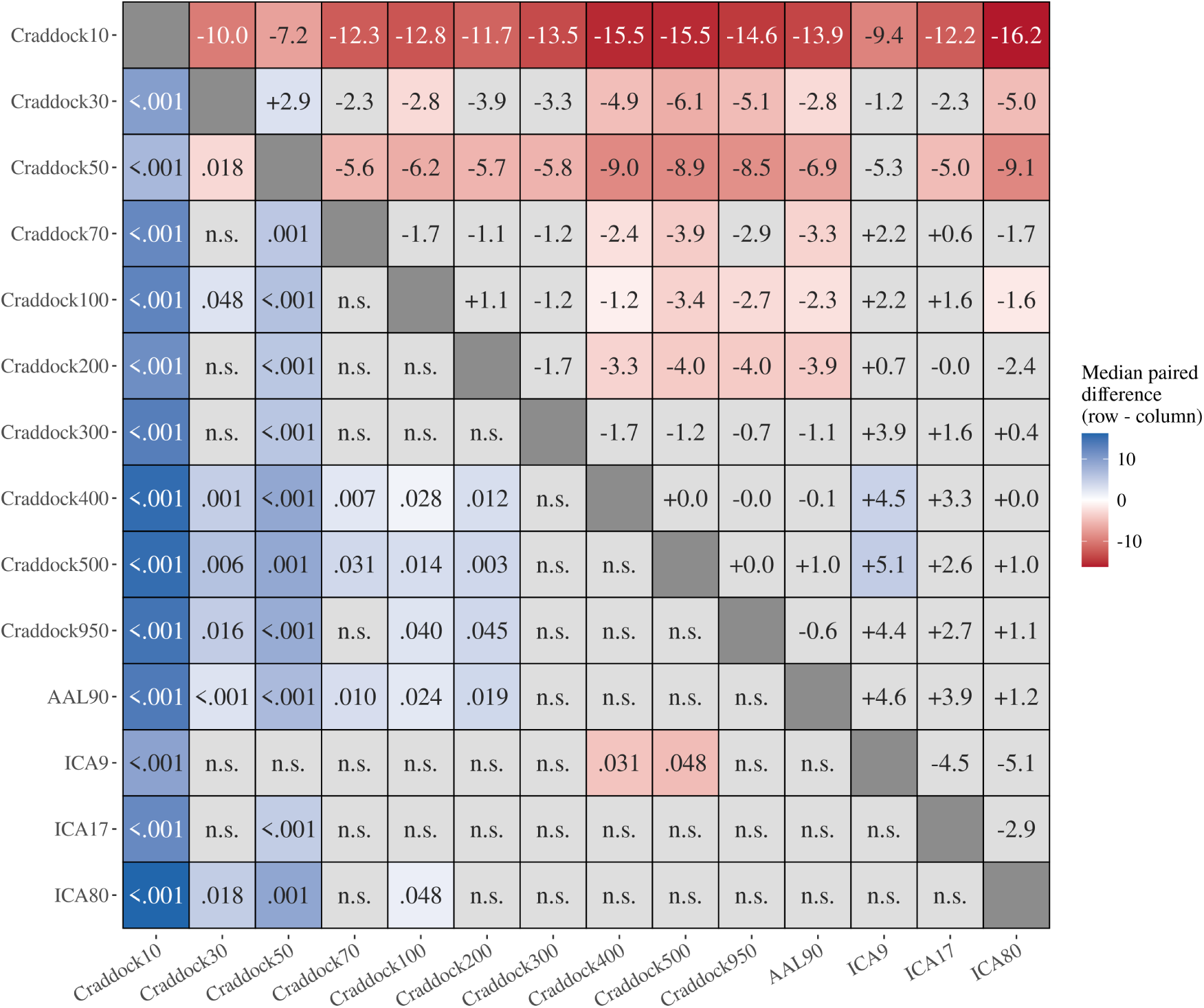
Pairwise comparisons between parcellations on the PCA-reduced features. Each cell is a paired Wilcoxon signed-rank test on the PCA-reduced, non-lagged results, matched on classifier (FDR-adjusted). Colour encodes the median paired difference in accuracy between the row and the column parcellation: blue where the row performed better, red where it performed worse. The matrix is symmetric: cells above the diagonal give the median difference, cells below it the adjusted *p* for the same pair, written without its leading zero. Non-significant cells are grey and labelled n.s. in place of the *p*. The Craddock levels above 300 regions are included, being run with PCA only.

**Figure S2:**
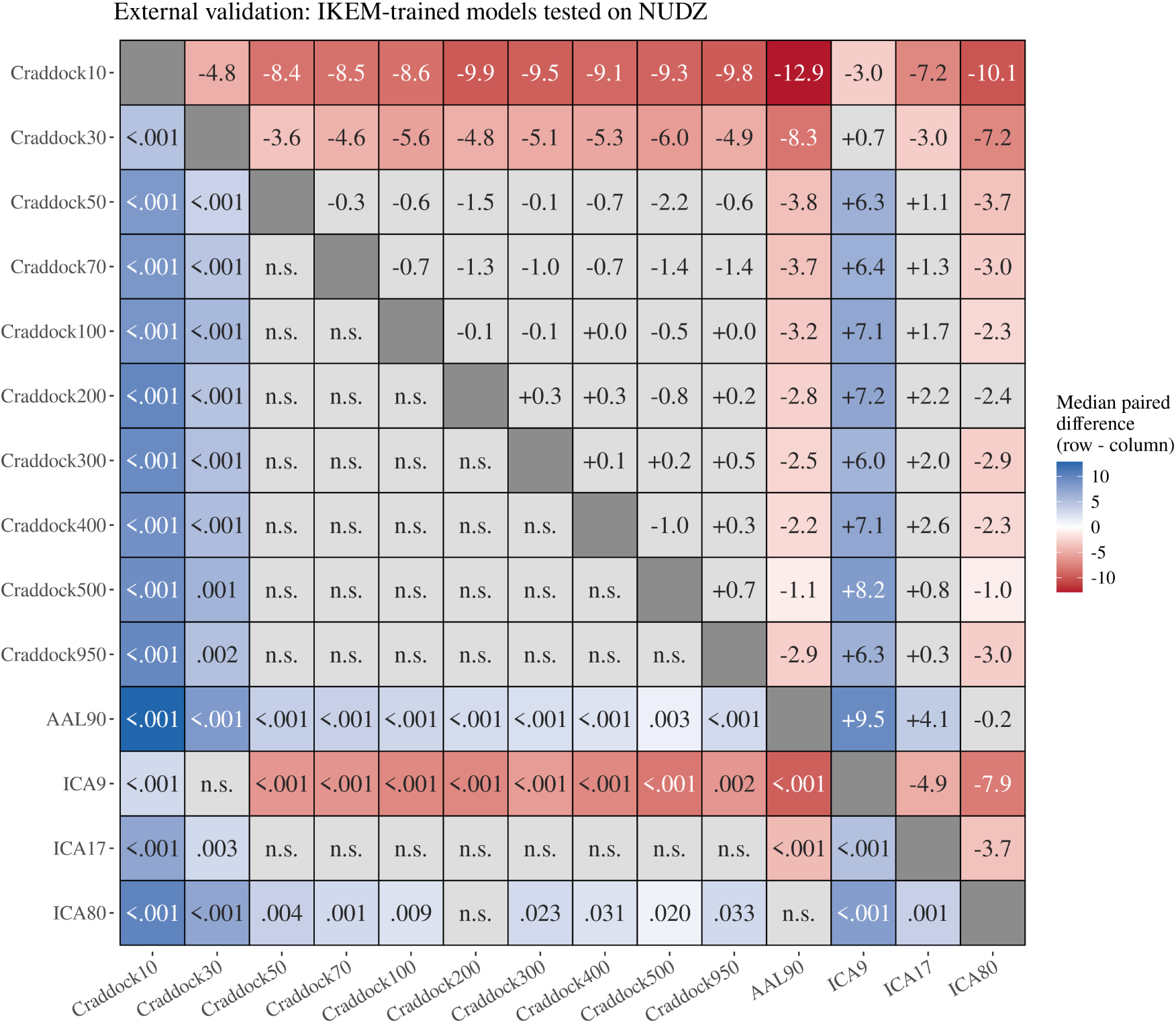
External validation (IKEM-trained models tested on NUDZ): pairwise comparisons between parcellations on the PCA-reduced features. Each cell is a paired Wilcoxon signed-rank test on the PCA-reduced, non-lagged results, matched on classifier (FDR-adjusted). Colour encodes the median paired difference in accuracy between the row and the column parcellation: blue where the row performed better, red where it performed worse. The matrix is symmetric: cells above the diagonal give the median difference, cells below it the adjusted *p* for the same pair, written without its leading zero. Non-significant cells are grey and labelled n.s. in place of the *p*. The Craddock levels above 300 regions are included, being run with PCA only.

**Figure S3:**
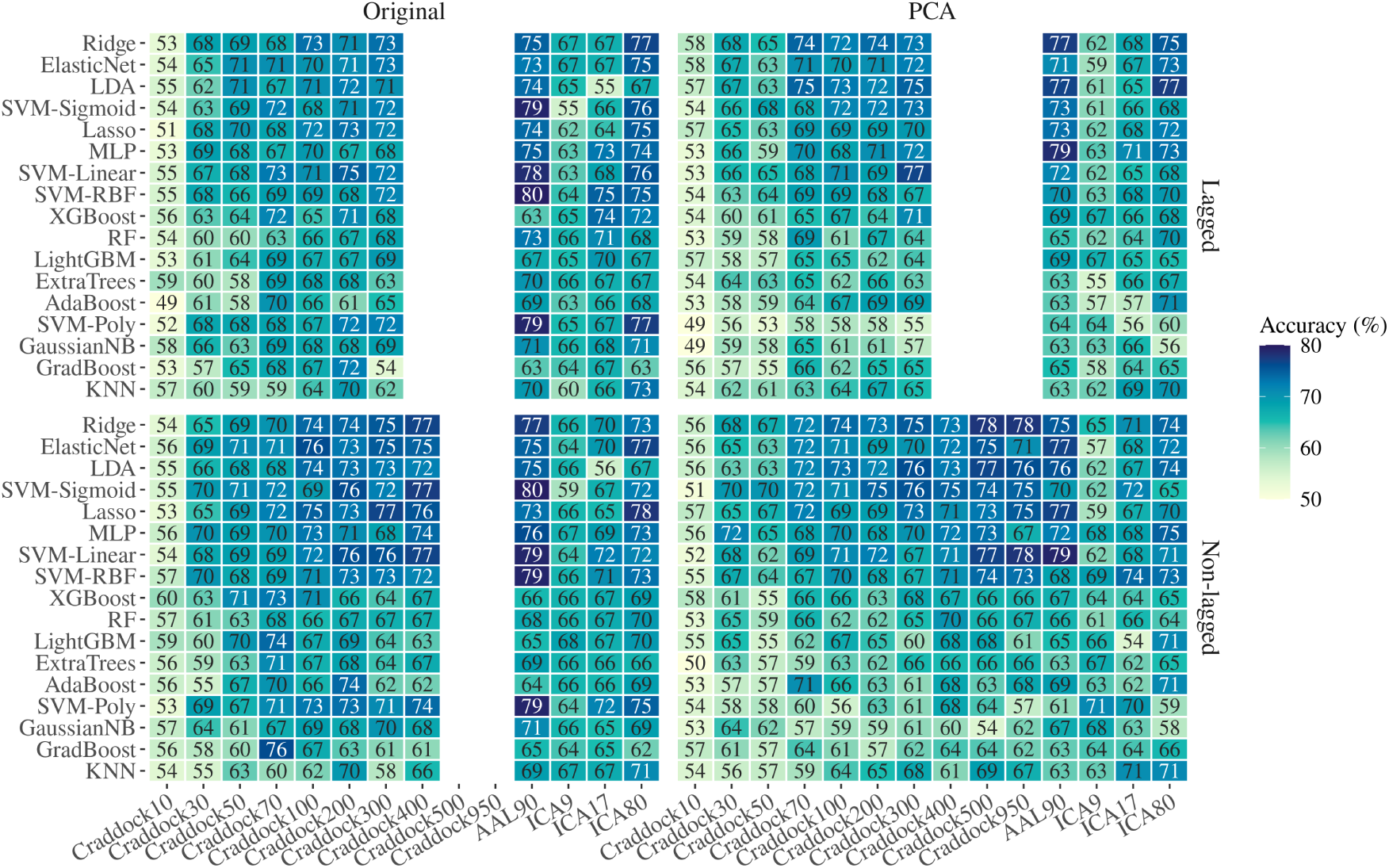
Accuracy of every classifier at every parcellation, split by connectivity type (rows) and by whether the features were PCA-reduced (columns). Each cell is the mean over the five outer folds, rounded to whole percentage points. Classifiers are ordered by their own median accuracy across parcellations, worst at the bottom. Fill encodes accuracy on a fixed 50–80% scale. Craddock400 was run non-lagged only, and Craddock500 and Craddock950 non-lagged and with PCA only.

**Figure S4:**
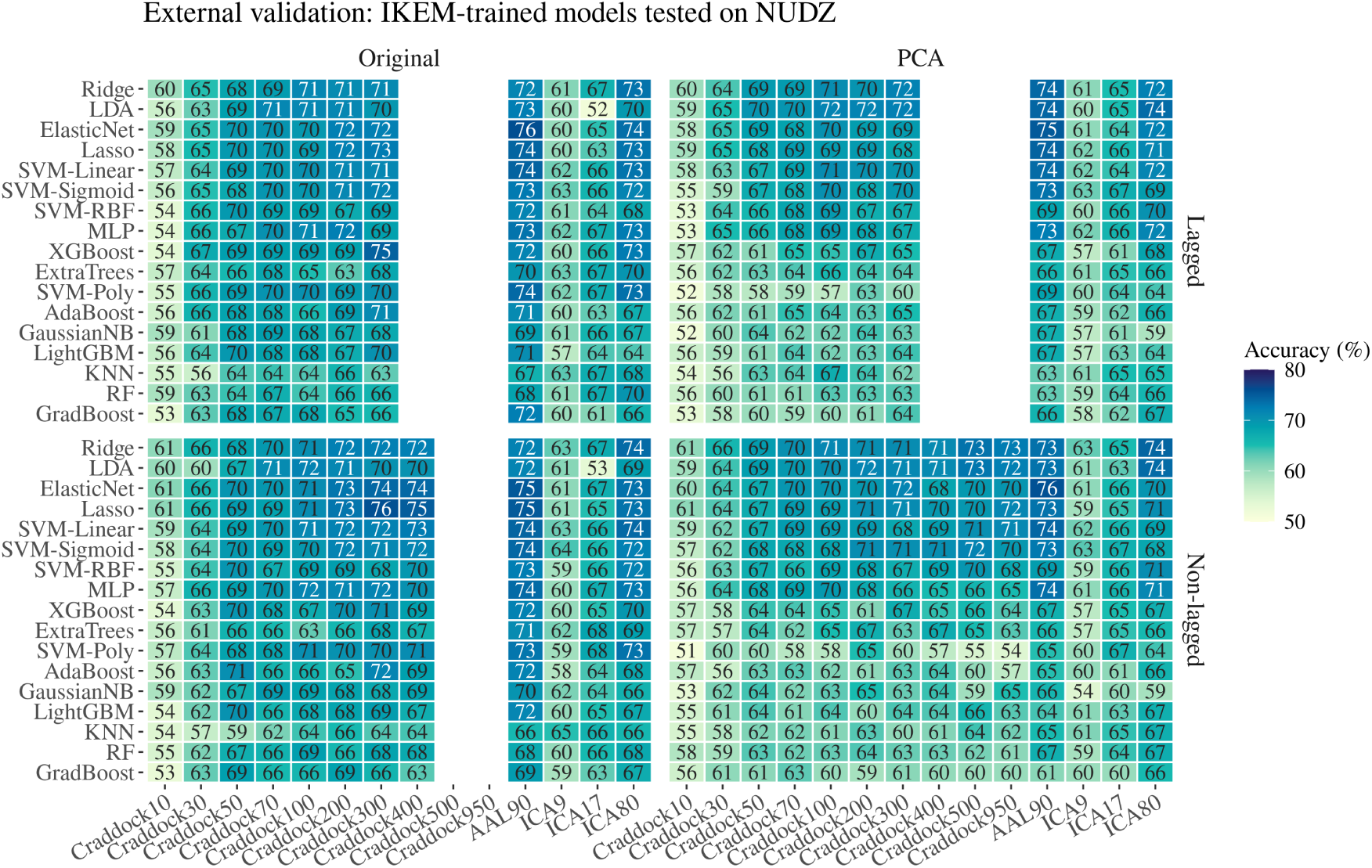
External validation (IKEM-trained models tested on NUDZ): accuracy of every classifier at every parcellation, split by connectivity type (rows) and by whether the features were PCA-reduced (columns). Each cell is the mean over the five IKEM-fold models, rounded to whole percentage points. Classifiers are ordered by their own median accuracy across parcellations, worst at the bottom. Fill encodes accuracy on a fixed 50– 80% scale. Craddock400 was run non-lagged only, and Craddock500 and Craddock950 non-lagged and with PCA only.

**Figure S5:**
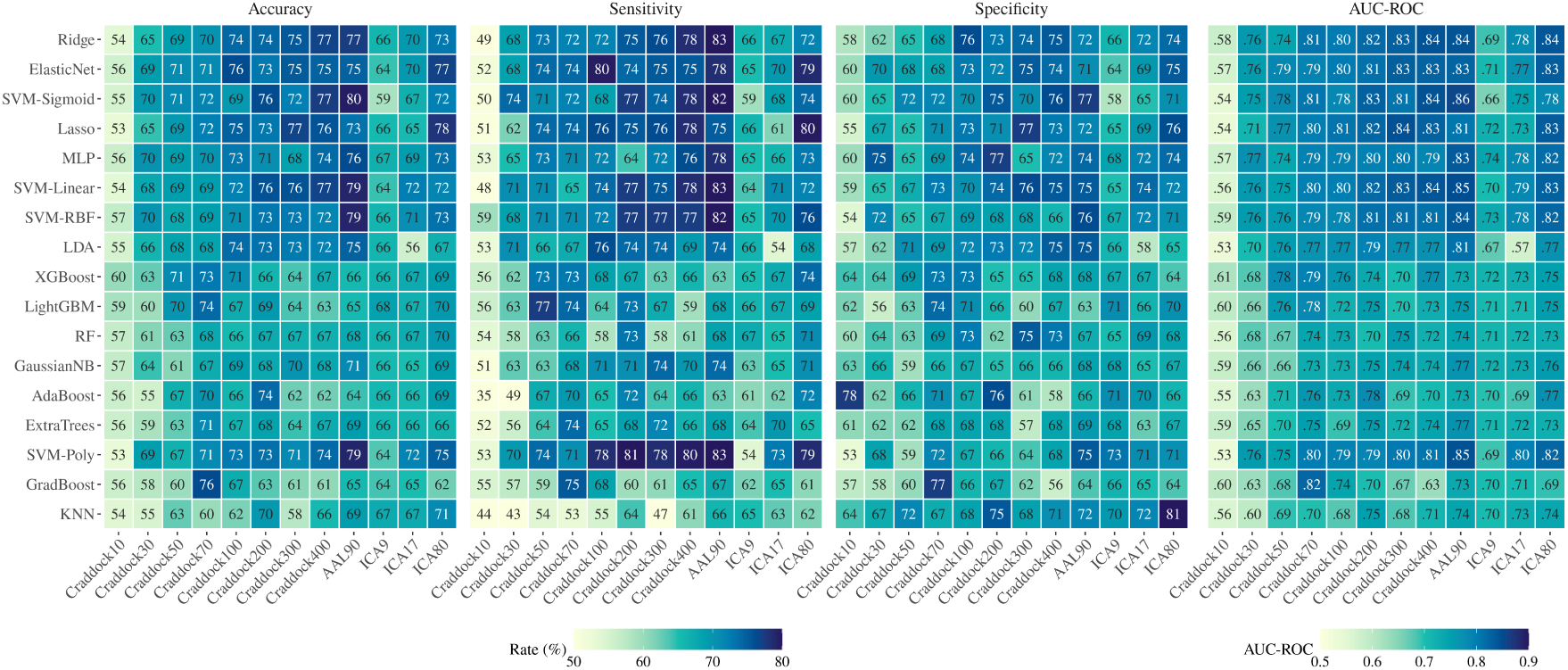
All four metrics for every classifier and parcellation; non-lagged connectivity, no PCA. Each cell is the mean over the five outer folds; the rates are rounded to whole percentage points and AUC-ROC to two decimals. The three rate metrics share a fixed 50–80% scale; AUC-ROC is drawn separately with its own scale.

**Figure S6:**
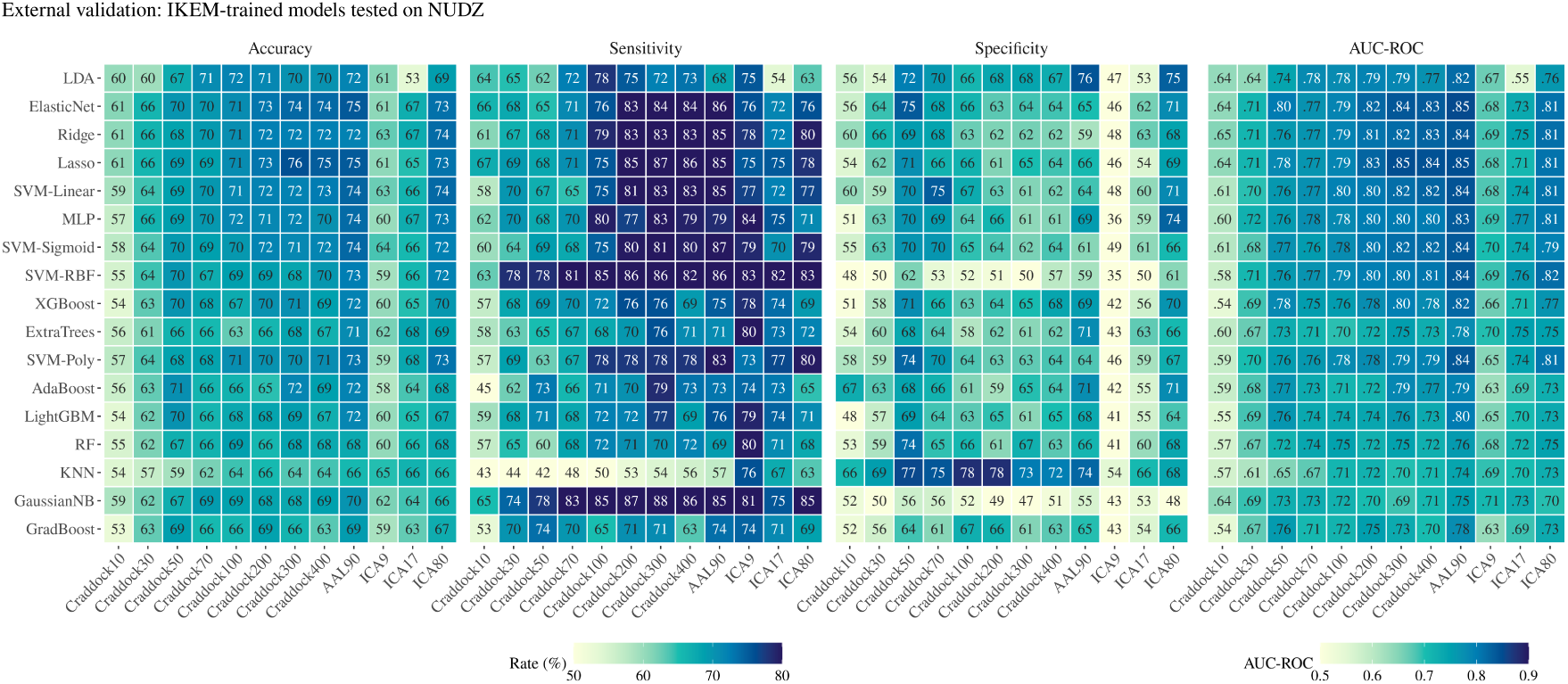
External validation (IKEM-trained models tested on NUDZ): all four metrics for every classifier and parcellation; non-lagged connectivity, no PCA. Each cell is the mean over the five IKEM-fold models; the rates are rounded to whole percentage points and AUC-ROC to two decimals. The three rate metrics share a fixed 50–80% scale; AUC-ROC is drawn separately with its own scale.

